# The unique coding sequence of *pmoCAB* operon from type Ia methanotrophs simultaneously optimizes transcription and translation

**DOI:** 10.1101/543546

**Authors:** Juan C. Villada, Maria F. Duran, Patrick K. H. Lee

## Abstract

Understanding the interplay between genotype and phenotype is a fundamental goal of functional genomics. Methane oxidation is a microbial phenotype with global-scale significance as part of the carbon biogeochemical cycle, and is a sink for greenhouse gas. Microorganisms that oxidize methane (methanotrophs) are taxonomically diverse and widespread around the globe. Recent reports have suggested that type Ia methanotrophs are the most prevalent methane-oxidizing bacteria in different environments. In methanotrophic bacteria, complete methane oxidation is encoded in four operons (*pmoCAB, mmoXYZBCD, mxaFI*, and *xoxF*), but how evolution has shaped these genes to execute methane oxidation remains poorly understood. Here, we used a genomic meta-analysis to investigate the coding sequences that encode methane oxidation. By analyzing isolate and metagenome-assembled genomes from phylogenetically and geographically diverse sources, we detected an anomalous nucleotide composition bias in the coding sequences of particulate methane monooxygenase genes (*pmoCAB*) from type Ia methanotrophs around the globe. We found that this was a highly conserved sequence that optimizes codon usage in order to maximize translation efficiency and accuracy, while minimizing the synthesis cost of transcripts and proteins. We show that among the seven types of methanotrophs, only type Ia methanotrophs possess a unique coding sequence of the *pmoCAB* operon that is under positive selection for optimal resource allocation and efficient synthesis of transcripts and proteins in environmental counter gradients with high oxygen and low methane concentrations. This adaptive trait possibly enables type Ia methanotrophs to respond robustly to fluctuating methane availability and explains their global prevalence.

## Introduction

Microbial methane oxidation plays a number of fundamental roles in the global ecosystem^1^. Methane-oxidizing microorganisms can mitigate methane emissions by acting as methane sinks^2,3^ and thereby reduce the contribution of methane to climate change^4^. Microbial oxidation of methane also provides an entry point for methane into the global food web, and can serve as a primary carbon source for large trophic systems^5,6^. Methane oxidation is a globally distributed phenotype expressed in microorganisms from diverse taxonomic groups. Based on a range of phenotypic and phylogenetic features, methanotrophic bacteria are categorized into seven major types^7^: Ia, Ib, Ic, IIa, IIb, III, and the candidate division NC10. The distribution of marker gene sequences for the major methanotroph types suggests they are differentially prevalent across environments^7^. In bacteria, methane oxidation involves two enzymatic steps where methane is first converted to methanol by a soluble (sMMO) or particulate (pMMO) methane monooxygenase, then methanol is oxidized to formaldehyde by a pyrroloquinoline quinone-containing methanol dehydrogenase that can be calcium-dependent (Mxa) or lanthanide-dependent (Xox). The resulting formaldehyde can be directed to energy production or biomass synthesis^8^. sMMO is encoded in the six-gene operon *mmoXYZBCD*, while pMMO is encoded in the three-gene operon *pmoCAB*. Of the methanol dehydrogenases, Mxa is encoded in the two-gene operon *mxaFI*, and Xox by the gene *xoxF*.

In evolutionary history, methane oxidation appeared at around the same time as oxygenic photosynthesis, nitrogen fixation, nitrification and denitrification^9^, and it is possible that the emergence of methanotrophy occurred soon after the last universal common ancestor^10^. Hence, evolution has likely shaped methanotrophs with many as yet undiscovered properties^10^. One unexplored question of fundamental importance to our understanding of methanotrophy is how the genes that encode the methane oxidation metabolic module have been shaped by evolution to efficiently execute methanotrophy. In the three cellular domains and in viruses, evolution has selected gene sequences that perform cellular functions beyond just encoding the amino acid composition of proteins. For example, gene sequences have nucleotide composition and codon usage that direct mRNA folding^11,12^, transcript abundance^13^, mRNA degradation^14^, RNA toxicity^15^, protein synthesis^16^, cotranslational protein folding^17^, and promote the interaction of peptides with the Signal Recognition Particle^18^. Gene sequences can also affect the cellular economy of protein synthesis^19^, reduce the metabolic burden of nucleotide synthesis by incorporating less expensive nucleotides^20^, and allocate resources required for transcription^21^ and translation^22^.

In this study, we performed an in-depth genomic meta-analysis to investigate the coding sequences of the genes that encode the methane oxidation metabolic module in bacteria. We found that evolution has shaped the *pmoCAB* operon of type Ia methanotrophs with a unique coding sequence that optimizes resource allocation by reducing the biosynthetic costs of transcription and translation while ensuring translation efficiency and accuracy. This study provides novel insights into the molecular biology and evolution of methanotrophic bacteria, and extends our understanding of the mechanisms developed by nature to sustain metabolism and life on Earth.

## Results

### Meta-analysis of methanotroph genomes

To investigate the coding sequence of genes encoding the methane oxidation metabolic module, we analyzed 67 methanotrophic bacterial genomes including 49 isolate genomes and 18 metagenome-assembled genomes (MAGs) (Additional file 1: Figure S1, Additional file 2). These genomes originated from different sampling locations around the globe (Additional file 1: Figure S1a) and were phylogenetically diverse (Additional file 1: Figure S1b), comprising 19 genera from five families (Additional file 2) and with GC contents ranging from 40% (*Methylacidiphilum kamchatkense* Kam1) to 65% (*Methylosinus trichosporium* OB3b). The 67 methanotroph genomes were categorized into the seven major types^7^: Ia (*n* = 36), Ib (*n* = 6), Ic (*n* = 3), IIa (*n* = 11), IIb (*n* = 5), III (*n* = 4), and NC10 (*n* = 2).

### Type Ia *pmoCAB* coding sequences have an anomalous nucleotide composition bias

We measured the distribution of nucleotide composition bias of all coding sequences in each genome (Additional file 1: Figure S1b). The guanine + cytosine (GC) content of every coding sequence was calculated, as well as the GC content in the third nucleotide of each codon (GC_3_). In genomes of methanotroph types Ia, Ib, IIa, IIb, and NC10, the GC_3_ content of coding sequences tended to be higher than the GC, while in type III genomes GC_3_ tended to be lower than GC (Additional file 1: Figure S1b). The distributions in type Ic genomes were variable, and the non-methanotrophic bacterium *Bacteroides ovatus*, included as an outgroup, had no difference between GC_3_ and GC.

Next, we analyzed the nucleotide composition bias of the 12 coding sequences in the four operons forming the methane oxidation metabolic module (Additional file 1: Figure S2). All coding sequences from all seven types showed patterns of relative GC/GC_3_ bias consistent with the whole genomes (Additional file 1: Figure S1), except for *pmoCAB* coding sequences of type Ia methanotrophs where GC_3_ content was lower than GC (Fig. 1). The *pxmABC* and *hao* coding sequences were selected as controls as they appear in most of the methanotroph genomes and are not part of the methane oxidation metabolic module. As expected, these coding sequences showed a pattern of relative GC/GC_3_ bias consistent with the whole genomes (Additional file 1: Figure S2). Similar to the type Ia *pmoCAB*, the type III coding sequences tended to have lower GC_3_ content compared to GC (Additional file 1: Figure S2), consistent with the whole-genome pattern in type III (Additional file 1: Figure S1).

**Fig. 1.**
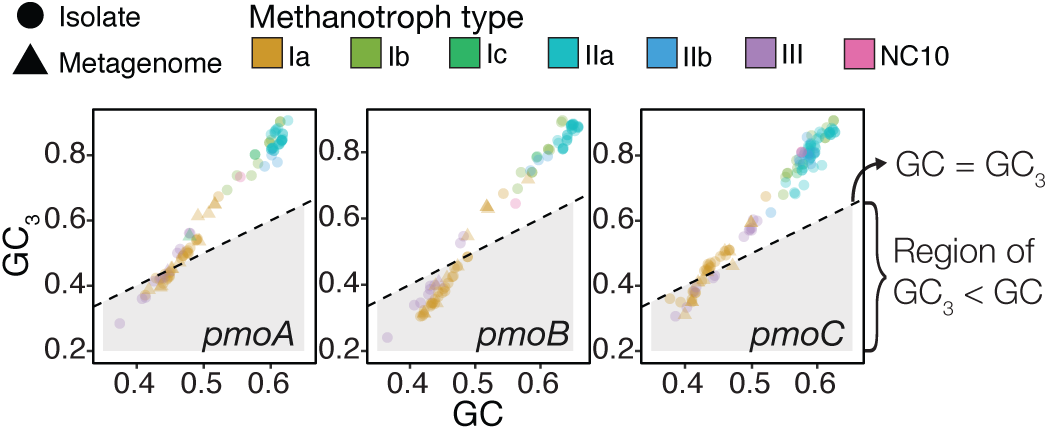
Nucleotide composition bias of *pmoCAB* coding sequences. An extended analysis of all coding sequences together with controls is shown in Additional file 1: Figure S2.

### The anomalous nucleotide composition of type Ia *pmoCAB* coding sequences provides an atypical codon usage bias

The anomalous nucleotide composition of type Ia *pmoCAB* coding sequences led us to hypothesize that this may generate an atypical bias in codon usage. To test whether the codon usage differs among coding sequences in the methane oxidation metabolic module of each methanotroph, we first calculated the relative synonymous codon usage (RSCU) in every coding sequence (controls were also included). On ordination of the RSCU values of type Ia coding sequences (Additional file 1: Figure S3a), *pmoCAB* coding sequences clustered separately (Euclidean distance PERMANOVA, F = 36.55, *p* < 0.01) and the effect of source operon in the separation was strong (*R*^2^ = 0.37). A reduced effect of source operon was observed in the clustering pattern for coding sequences of types Ib (F = 3.82, *p* < 0.01, *R*^2^ = 0.21) and IIa (F = 12.67, *p* < 0.01, *R*^2^ = 0.24), while weak effects (*R*^2^ < 0.2) or non-significant probabilities (*p* ≥ 0.01) were observed for types Ic, IIb, III and NC10 (Additional file 1: Figure S3).

This result indicates that the type Ia *pmoCAB* coding sequences have an atypical codon usage bias compared to other coding sequences in the methanotrophy module. We hypothesized that this may be an adaptation to allow for higher expression of these genes. To test this, we calculated a Codon Adaptation Index (CAI) that compared the codon usage of a coding sequence to the codon usage of a reference set of sequences from the same genome. The first CAI, defined as CAI_*genome*_, used the codon frequency of all coding sequences in the genome as a reference, while the second CAI, defined as CAI_*ribosome*_, used the codon frequency of only ribosomal protein genes as a reference. CAI_*ribosome*_ thus served as a proxy to represent the codon usage of highly expressed genes. The comparison between the CAI_*genome*_ and CAI_*ribosome*_ showed that *mxaF* coding sequences tended to have similar codon usage to both reference sets in all types of methanotrophs (Additional file 1: Figure S2a), which was also observed for *mxaI* and *xoxF* coding sequences in types Ib and Ic. Type Ia *mxaI* coding sequences tended to deviate from the codon bias of ribosomal protein genes and towards the genome-scale codon usage pattern, while the opposite was observed for type IIa and IIb *mxaI* coding sequences. Type Ia *xoxF* coding sequences appeared in the top percentile ranks of CAI_*ribosome*_ and were less similar to the genomic codon usage. A similar but much stronger pattern was observed in type Ia *pmoCAB* coding sequences where they appeared in the bottom 10% of CAI_*genome*_ values, but at much higher percentile ranks for CAI_*ribosome*_ (Additional file 1: Figure S2a). The results for all coding sequences in the methane oxidation module and controls are given in Additional file 1: Figure S4a. Overall, the extreme deviation of type Ia *pmoCAB* coding sequences from the genomic norm shows that they have a codon usage more similar to that of the ribosomal protein-coding sequences.

This deviation of codon usage from the genomic signature could be due to a higher codon usage bias in *pmoCAB* coding sequences. To test this, we computed the effective number of codons (ENC) of every coding sequence in all type Ia methanotrophs. ENC measures the number of synonymous codons used to encode amino acids. Thus, the ENC is 61 if a coding sequence uses all codons equally to produce all twenty standard amino acids, whereas the ENC is 20 if only one codon is used per amino acid (i.e. a high codon usage bias). Contrary to our hypothesis, the ENC of type Ia *pmoCAB* coding sequences was not significantly different (two-tailed Wilcoxon signed rank-test, 99% confidence level) to that of *mxaFI* and *xoxF* coding sequences (Additional file 1: Figure S2b), while the ENC of *pmoCAB* coding sequences was significantly lower than coding sequences of ribosomal proteins and whole genomes (Additional file 1: Figure S4b). Overall, this indicates that even though the codons in coding sequences of *pmoCAB* and ribosomal proteins tend to be similar (Fig. 2a), the codon usage bias of *pmoCAB* coding sequences is higher. We hypothesized that if the type Ia *pmoCAB* coding sequences have a codon usage bias similar to other coding sequences in the methane oxidation metabolic module (Fig. 2b) but their codon usage frequencies are divergent (Additional file 1: Figure S2a), there should be a clear association between the level of codon bias (Fig. 2b) and the anomalous nucleotide composition bias (Fig. 1). We found that type Ia *pmoCAB* coding sequences increase their level of codon bias at the expense of decreasing the GC_3_ content (Fig. 2c), which contrasts with all other methanotroph types and coding sequences tested (Additional file 1: Figure S5). The *pmoCAB* coding sequences of type III show similar patterns to type Ia, but this is expected based on the whole-genome nucleotide bias (Additional file 1: Figure S1).

**Fig. 2.**
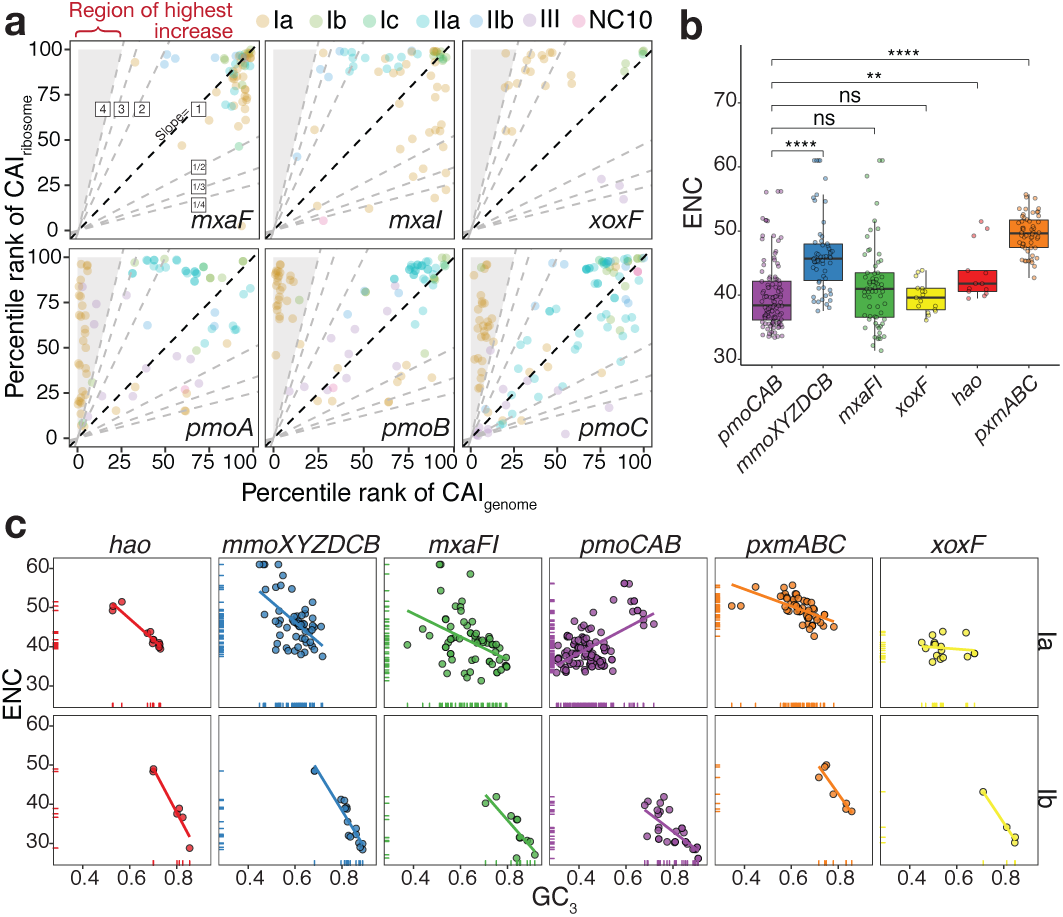
Analysis of the codon usage bias. **a** Analysis of the Codon Adaptation Index (CAI). CAI indexes were converted to percentile ranks based on the relative distribution of CAI in each methanotroph. Dashed lines with varying slopes delineate the variations between the percentile ranks of both indexes. **b** Analysis of the distribution of effective number of codons (ENC) values of coding sequences from type Ia methanotrophs. The two-tailed Wilcoxon signed rank-test was applied to test the difference of ENC values between the *pmoCAB* and other coding sequences. Non-significant result of the tests is denoted by *ns* (*p* > 0.01); \*\**p* ≤ 0.01; \*\*\**p ≤* 0.001; \*\*\*\**p* ≤ 0.0001. **c** Analysis of the ENC as a function of GC_3_ with the line representing the fit of a linear model between ENC and GC_3_.

### Codon biases in methanotrophy operons optimize translation efficiency

The anomalous nucleotide composition bias of type Ia *pmoCAB* coding sequences suggests that there is a fitness advantage to either the particular codons selected or to having a lower codon diversity. We hypothesized that this advantage may lie in increased translation efficiency. To evaluate this, we first analyzed the encoded tRNA pool of all isolate genomes. MAGs were not included in this analysis as their tRNA pool could be partial due to incomplete genome assembly. We found that the main difference among methanotrophs is that the type Ia genomes lack tRNAs with cytosine in the 3^*1*^-end of the anticodon (Additional file 1: Figure S6). Taking wobble-pairing into account, we queried whether the codon usage bias is an optimization strategy in which type Ia *pmoCAB* codon usage has been selected to better match available tRNA isoacceptors and thereby improve translation efficiency by recruiting the most abundant tRNAs. To address this question, we estimated the translation efficiency of coding sequences using the tRNA adaptation index (tAI). The tAI considers exact and wobble-pairing to estimate how efficiently a given coding sequence would be translated based on its codon usage and the genomic tRNA pool. We found that even though the preference for individual codons is different between type Ia *pmoCAB, mxaF*, and *xoxF* coding sequences, their translation efficiency is similarly ranked around the top 10^*th*^ percentile in each genome; the only coding sequence showing significant difference to *pmoC* is *mxaI* (*t*-test, 99% confidence level) (Fig. 3a). We included a control analysis to test the difference between the *pmoC* coding sequence and controls (Fig. 3a) and found that in this case the percentile rank of tAI is significantly different (*t*-test, 99% confidence level). The tAI results of other coding sequences and methanotrophs can be found in Additional file 1: Figure S7.

**Fig. 3.**
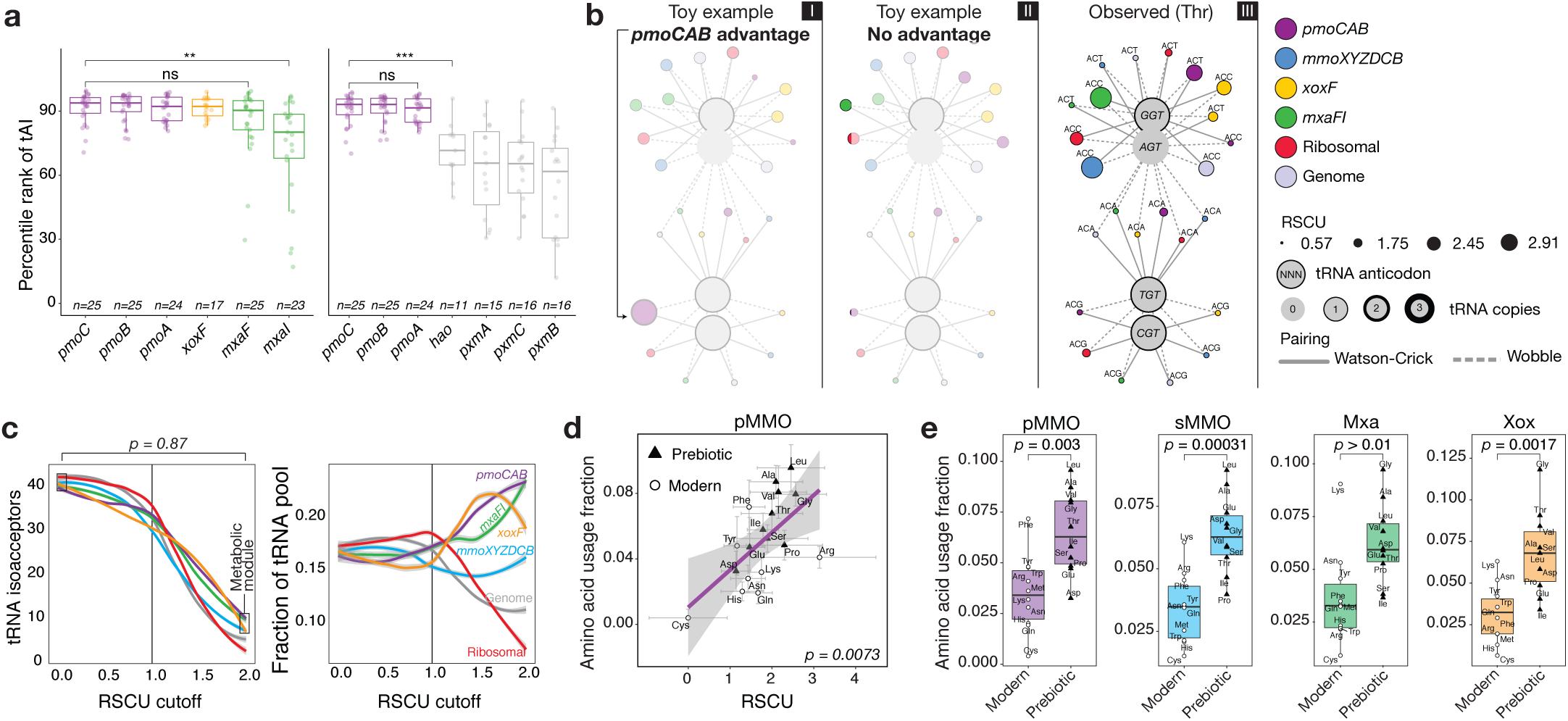
Strategies for optimal translation. **a** The tRNA adaptation index (tAI) was used to estimate the translation efficiency of coding sequences. tAI values were converted to percentile ranks based on the relative distribution of tAI in each methanotroph. Coding sequences were ordered by the mean of the tAI percentile rank and the two-tailed *t*-test was applied to test whether the percentile rank of tAI is significantly different between the *pmoC* coding sequence (highest mean) and other coding sequence. *ns* (*p* > 0.01); \*\*\**p ≤* 0.001. **b** Network of the interaction between codons and tRNA in type Ia methanotrophs. The integrated network was composed by median RSCU values, median number of tRNA copies, and the codon-anticodon pairing rules. The two toy examples illustrate the expected networks when *pmoCAB* coding sequences have a codon usage bias that (I) grants an advantage (marked by an arrow) in the competition for the tRNA pool or (II) does not grant any advantage. The observed network for threonine is shown in (III) and all other amino acids are shown in Additional file 1: Figure S11. **c** Quantitative analysis of the codon-tRNA interaction network. The figure shows how many copies of tRNA a gene can access according to a RSCU cut-off varying from 0.0 to 2.0 with a step size of 0.01. The reference line at RSCU = 1.0 indicates the change from the region of non-preference codon usage (RSCU < 1.0) to the region of high-preference for individual codons (RSCU > 1.0). The colored lines show the fit of a polynomial surface using local fitting. The Chi-squared test of proportions was applied to test whether the proportion of tRNAs available to coding sequences in the methane oxidation metabolic module is different between RSCU = 0 and RSCU = 2. **d** The median usage of each amino acid in each translated coding sequence as a function of the median RSCU value of the codon exhibiting the highest preference for each amino acid for type Ia methanotrophs. The *p*-value corresponds to the significance of the linear regression. **e** Usage of prebiotic (cheap) and modern (expensive) amino acids in the proteins of the methane oxidation metabolic module of type Ia methanotrophs. The *t*-test was used to evaluate the difference between the two groups.

Overall, these results suggest that for type Ia methanotrophs, the translation efficiency of *pmoCAB, mxaF*, and *xoxF* coding sequences are similar and these coding sequences have been optimized in parallel through different codon usage biases. A possible evolutionary explanation for these observations is that the distinct codon usage bias of each coding sequence causes these genes to segregate between different tRNA isoacceptors, allowing simultaneous translation of the coding sequences required for methane oxidation while avoiding competition for the same tRNAs. To investigate this, we reconstructed the protein synthesis network accounting for the distribution of tRNAs among coding sequences of the metabolic module (Fig. 3b). We also included in the network the median RSCU value of each codon per coding sequence of type Ia methanotrophs (Additional file 1: Figure S8-S9), so that we can account for the codon preference and its interaction with tRNAs through different pairing mechanisms. The median codon usage biases of the whole genome and coding sequences of ribosomal proteins (*n* = 1,246, median ribosomal proteins per genome = 55, standard deviation = 3.75) were integrated in the network to compare the links between codons and tRNAs outside the methane oxidation module (details on the tRNA pool of each genome are given in Additional file 1: Figure S10). The resulting networks (Fig. 3b; see Additional file 1: Figure S11 for networks for all amino acids) did not support the hypothesis that the codons favored by each operon provided differential access to tRNA isoacceptors. We also analyzed how access to different tRNA isoacceptors changes with variations in individual codon preference of coding sequences (Fig. 3c). We found that the different codon usage biases do not provide the operons with access to significantly different fractions of the tRNA pool (Chi-square test for equality of proportions, *p* = 0.87). Taken together, these results suggest that the codon usage bias of type Ia *pmoCAB* coding sequences does not provide a fitness advantage either in translation efficiency or in differential access to different tRNA isoacceptors when compared against the *mxaF* and *xoxF* coding sequences. Despite the differences in codon usage between operons from the methane metabolic module, simultaneous translation of these operons would still require competition for the same tRNA isoacceptors.

### Type Ia *pmoCAB* coding sequences optimize translation accuracy

We hypothesized that the divergent codon bias of *pmoCAB* coding sequences could be a strategy to optimize translation accuracy. This strategy prevents the incorporation of incorrect amino acids in the nascent protein^23,24^ by reducing the binding probability of near-cognate tRNA isoacceptors (i.e. tRNAs that differ by one base from the correct anticodon) for the ribosomal A-site^25^. Such an optimization for accuracy can be achieved at the coding sequence level through specialization of the codon-anticodon interaction in which an amino acid is encoded by a frequent and optimal codon that is recognized by an abundant cognate tRNA^26^. We hypothesized that if translation accuracy is a selective pressure acting on a coding sequence, the positive selection for an optimal codon (or the purifying selection acting on non-optimal codons) should increase as that codon’s amino acid is increasingly used in the encoded protein.

To test this, we first calculated the amino acid usage of each protein (Additional file 1: Figure S12) and mapped the RSCU with the highest median per amino acid (Additional file 1: Figure S8) to the median frequency of amino acid usage. Finally, we used a linear regression to analyze the relationship between the codon preference in coding sequences with the amino acid usage frequency in their proteins. Consistent with our translational accuracy hypothesis, we found that among all methanotrophs and all coding sequences of the methane oxidation metabolic module, only the type Ia *pmoCAB* coding sequences showed a significant linear relationship (*p* < 0.01, Fig. 3d, Additional file 1: Figure S13). This is especially notable for the two amino acids with the highest codon degeneracy (i.e. the 6-fold Leu and Arg).

### Minimizing protein synthesis cost through favoring of prebiotic amino acids

The use of CAI, ENC, RSCU and tAI enabled us to investigate the role of synonymous codons in optimization for translation efficiency and accuracy. However, as these measures account for amino acid usage biases, optimization strategies at the protein sequence level can be masked. We hypothesized that if translation optimization at the protein sequence level is achieved through reducing the synthesis cost of proteins^27,28^, there should be an increased usage of the prebiotic (cheaper) amino acids^29^ in the type Ia pMMO sequence. To test this, we first visually inspected the occurrence of prebiotic and modern amino acids in the linear regression between RSCU and amino acid usage (Fig. 3d), then quantitatively compared the median composition of cheaper and expensive amino acids in each protein (Fig. 3e). We observed that usage of prebiotic amino acids is significantly higher than modern (expensive) amino acids in pMMO, sMMO and Xox, while Mxa showed a non-significant difference (*t*-test, 99% confidence level; Fig. 3e).

### The codon usage of pmoCAB coding sequences encodes an effective resource allocation strategy to synthesize highly demanded transcripts

While the codon usage and nucleotide composition biases of coding sequences in the type Ia methane oxidation module does not seem to provide an advantage in translation efficiency, it is possible that they provide an efficiency advantage at the level of transcription. Previous reports on methanotrophic species^30–32^ and communities^33–35^ have shown that *pmoCAB* coding sequences are consistently highly expressed regardless of the experimental conditions, and the demand on cellular resources to transcribe *pmoCAB* coding sequences could be significant. To test whether the biases confer increased transcriptional efficiency, the transcriptomic datasets of three different type Ia methanotrophs obtained under different experimental conditions were examined. For illustration purposes, the results of *Methylobacter tundripaludum* 31/32 grown with methane as sole carbon source and supplemented with lanthanum^32^ are described here. Detailed results of the other methanotrophs and experimental conditions can be found in Additional file 1: Figure S14-S16.

To test the transcription optimization hypothesis, we first analyzed the purine and pyrimidine content of coding sequences in each genome and found that purine-derived residues occurred more frequently (52%) than pyrimidine nucleotides (Additional file 1: Figure S14a). We postulated that, if *pmoCAB* coding sequences are optimized for efficient transcription, they should have a higher pyrimidine than purine content, which reduces the mRNA synthesis cost as pyrimidines have fewer carbon and nitrogen atoms per molecule (Additional file 1: Figure S14a). We found that the *pmoCAB* were the most expressed transcripts and they have a higher pyrimidine than purine content (Additional file 1: Figure S14b), contrasting the *xoxF* and *mxaFI* coding sequences (Additional file 1: Figure S14b) and the genome-scale frequency (Additional file 1: Figure S14a). We calculated the mean pyrimidine content of 1,000 random combinations of three coding sequences Additional file 1: Figure S14c) and found that < 3.5% of the combinations had pyrimidine contents higher than the *pmoCAB* coding sequences.

To understand the effect of *pmoCAB* nucleotide frequency bias on cellular ribonucleotide composition, we analyzed the effect on the total ribonucleotide composition if we removed all the transcripts of an individual operon from the transcriptome. We calculated the total ribonucleotide composition by summing the ribonucleotide composition of all transcribed coding sequences in the transcriptome. We found that removing *mxaFI* and *xoxF* transcripts had negligible effect on the total ribonucleotide composition (Additional file 1: Figure S14d-f). However, removing *pmoCAB* transcripts caused a shift of 1% in the purine/pyrimidine balance, a shift of > 2% in the GC content, and a shift of 2% in uracil content (Additional file 1: Figure S14d-f).

If the nucleotide composition bias of *pmoCAB* coding sequences is an adaptation to reduce transcript synthesis cost, it would be expected that the atomic element investment per codon is reduced in *pmoCAB* transcribed coding sequences compared to others in the methane oxidation metabolic module and to the whole-genome average. To evaluate this, we calculated the atomic element composition of each transcribed coding sequence in the metabolic module and of all transcribed coding sequences in the genome based on their adenine, guanine, cytosine and uracil composition. We found that *pmoCAB* transcripts have fewer carbon, hydrogen, and nitrogen atoms per codon compared to other transcripts from the methane metabolic module and the genome-scale mean (Fig. 4a and 4b). However, although the codon usage of *pmoCAB* transcripts requires reduced investment of carbon, nitrogen, and hydrogen, this comes at the expense of increased oxygen demand, with the oxygen demand exceeding the genome-scale mean (Fig. 4b). We selected 1,000 random combinations of three transcribed coding sequences from the whole genome, calculated the per-codon element compositions, and compared them to the *pmoCAB* transcribed coding sequences demand (Fig. 4a and 4b). We found that < 3% of the combinations of transcribed coding sequences yielded a mean element composition below that of the *pmoCAB* transcribed coding sequences for nitrogen, carbon and hydrogen or above for oxygen. This pattern was even more pronounced for *Methylomicrobium alcaliphilum* 20Z (< 0.4%; Additional file 1: Figure S15g-h) and *Methylomicrobium buryatense* 5G (< 1.5%; Additional file 1: Figure S16g-h).

**Fig. 4.**
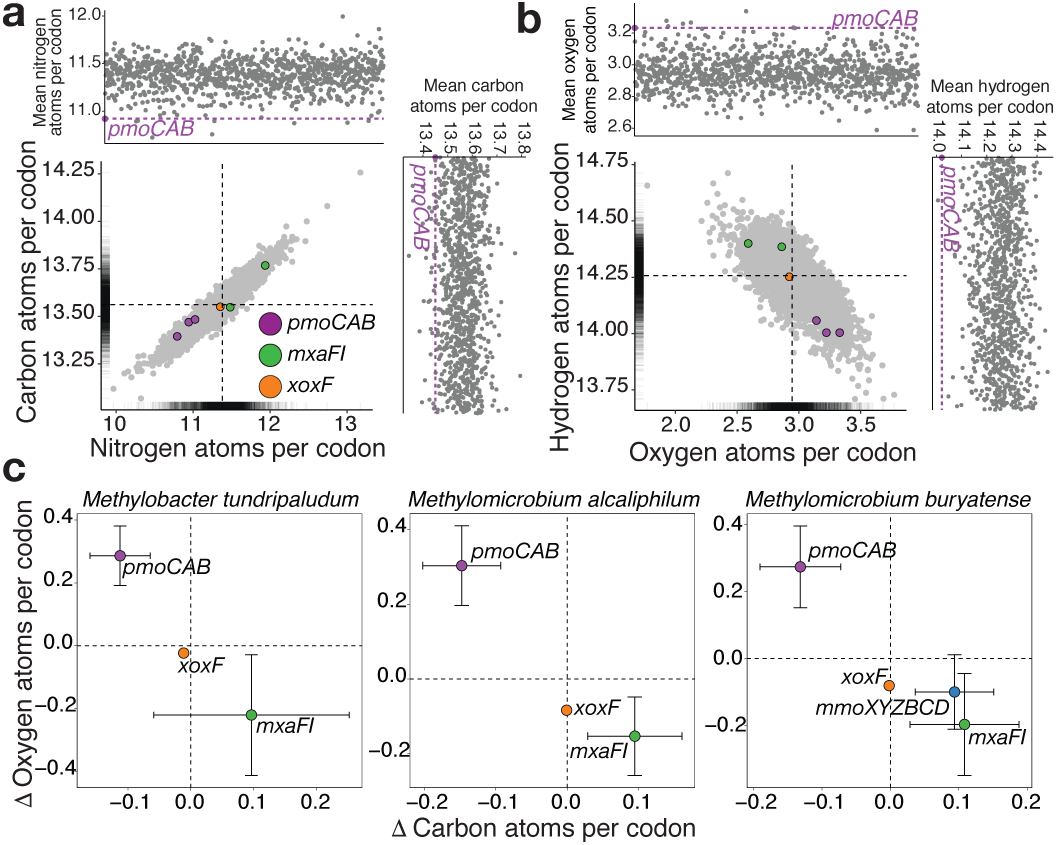
Strategies for optimal transcription in type Ia methanotrophs. **a, b** Elemental composition of transcribed coding sequences in *M. tundripaludum* 31/32. The main panel shows the number of element atoms per codon in each transcribed coding sequence with the dashed lines representing the mean of all transcribed coding sequences in the genome. The top and right panels show the mean number of element atoms per codon of the *pmoCAB* transcribed coding sequences and the mean number of element atoms per codon of 1,000 random combinations of three coding sequences. **c** A summary of the carbon and oxygen atom demands per codon of transcribed coding sequences. The difference (Δ) was calculated using the genomescale mean demand as the reference (dashed line). *M. buryatense* 5G possesses genes encoding both the pMMO and sMMO. An extended analysis for each methanotroph is shown in Additional file 1: Figure S14, S15 and S16.

Based on these results, an intriguing new question emerged: is the element composition of type Ia *pmoCAB* transcripts a reflection of the low methane and high oxygen composition of habitats where these methanotrophs are commonly found^10^? To assess this question, we calculated the distance (Δ) between the mean carbon and oxygen atom counts per codon for methane oxidation metabolic module transcripts and all transcripts from each genome. We found that the *pmoCAB* transcripts are the only set of transcripts in the methane oxidation metabolic module that had low carbon and high oxygen composition per codon (Fig. 4c). Finally, we assessed whether the elemental composition of the *pmoCAB* operon is a result of a genome-wide trend for reducing the atoms per codon as a function of mRNA abundance. Rejecting this hypothesis, we found only weak correlations between elemental content and mRNA abundance (Additional file 1: Figure S14i, S15i and S16i). Together, these results suggest an optimization strategy unique to type Ia *pmoCAB* coding sequences whereby highly demanded transcripts use fewer carbon, nitrogen, and hydrogen atoms at the expense of more oxygen atoms per codon. Given the high oxygen requirement for synthesis of *pmoCAB* transcripts, we hypothesized that type Ia genomes may harbor a genetic regulatory mechanism adjusting *pmoCAB* transcription in response to oxygen availability. Supporting this, we identified proteins containing the PAS domain, which can potentially act as oxygen sensors^36^, located near the *pmoCAB* coding sequences in the type Ia genomes of genera *Methylosarcina, Methylomonas, Methylomicrobium, Methylomarinum* (about ten coding sequences upstream), and *Methylobacter tundripaludum* 31/32 (five coding sequences downstream) (Additional file 1: Figure S17).

## Discussion

This study provides genomic evidence that the *pmoCAB* coding sequences of type Ia methanotrophs possess a unique adaptive trait manifesting as a strong nucleotide bias (Fig. 1) that fine-tune codon usage (Fig. 2) and can optimize methane oxidation through maximizing translation efficiency and accuracy (Fig. 3), while minimizing protein (Fig. 3) and transcript synthesis costs (Fig. 4). The discovery of this unique coding sequence was enabled by meta-analysis of a large number of isolate genomes and MAGs from diverse taxonomy and geography (Additional file 1: Figure S1). This finding illustrates a sophisticated adaptive linkage between molecular genotype and phenotype for methane oxidation.

From an environmental point of view, this adaptive trait would enable type Ia methanotrophs to be competitive and efficient in oxidizing methane, which is a gas with limited solubility and mass transfer in liquid^37^ and often found at low concentrations in the aerobic zone of the oxygen/methane countergradient^10^. The high oxygen and low carbon demands of the type Ia *pmoCAB* transcribed coding sequences (Fig. 4), relative to other transcribed coding sequences in the methane oxidation metabolic module and the rest of the transcriptome, reflect the efficiency of these strictly aerobic^38^ methanotrophs in colonizing and performing methane oxidation in environmental zones with high oxygen and low methane concentrations^35,39,40^ (Fig. 5a). Past reports on the molecular optimization of genetic sequences have shown that transcribed regions of genes can have fine-tuned nucleotide compositions that reduce their biosynthetic cost^20^. For example, DNA coding sequences are found to reduce nitrogen atoms per codon when nitrogen is scarce in the environment^41^, and a large proportion of bacterial genomes present a dual optimization strategy to reduce per-codon nitrogen demand while increasing translation efficiency^42^. Extending these findings, we show that type Ia *pmoCAB* coding sequences possess an adaptation that minimizes not only nitrogen but also carbon and hydrogen per transcribed codon, while reducing the metabolic burden of erroneous biosynthesis^25^ through optimal translation efficiency and accuracy. This optimization strategy likely increases the fitness of type Ia methanotrophs in the competition for methane and contributes to the prevalence of type Ia methanotrophs across diverse environments^7^. However, the transcript optimization strategy found in type Ia *pmoCAB* coding sequences results in an increased demand for oxygen atoms per codon. The high demand for oxygen to synthesize *pmoCAB* transcripts could be part of a mechanism to modulate cellular metabolism based on oxygen availability in the environment and to tightly regulate the first enzymatic step of the methane oxidation metabolic module at the transcription level. This is supported by the presence of genes encoding the PAS domain proteins, potentially acting as oxygen sensors^36^, flanking the *pmoCAB* coding sequences in the genomes of several type Ia methanotrophs (Additional file 1: Figure S17).

**Fig. 5.**
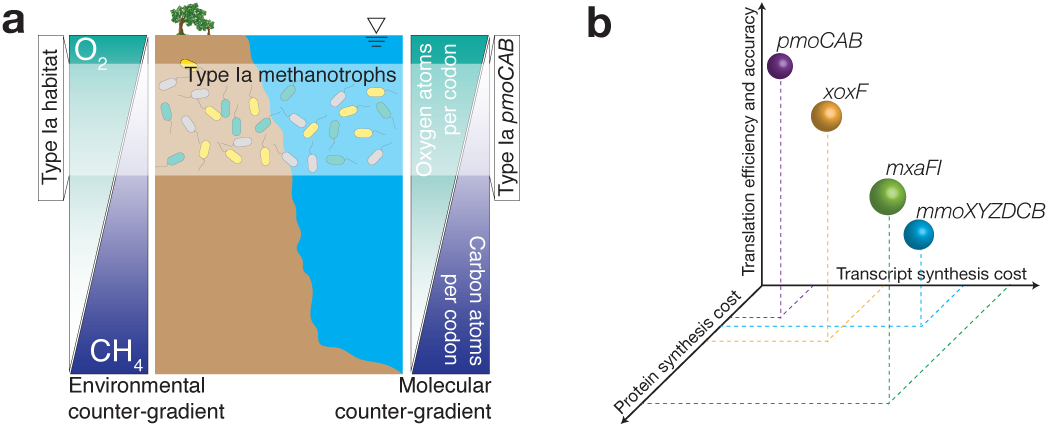
Schematic representation of the molecular characteristics of type Ia methanotrophs. **a** A cartoon depicting the environmental counter-gradient of oxygen and methane in terrestrial and aquatic settings and the potential coupling with the molecular demands of oxygen and carbon atoms per codon of the type Ia *pmoCAB* coding sequences. The demands are relative to the genome-scale mean. **b** The three levels of molecular optimization encoded in the coding sequences of the methane oxidation metabolic module of type Ia methanotrophs.

In the second enzymatic step of the methane oxidation metabolic module, methanol is oxidized to formaldehyde by the methanol dehydrogenase encoded by either the *mxaFI* or *xoxF* genes. We show here that the type Ia *pmoCAB, mxaFI*, and *xoxF* coding sequences have similar translation efficiency (Fig. 3a), similar access to tRNA isoacceptors (Fig. 3b and 3c), and a similar level of codon bias (Fig. 2b); however, their preference for individual codons (Additional file 1: Figure S3a) and translation accuracy (Fig. 3d) is different. Considering only the genes for the second enzymatic step, the difference in codon usage between the *mxaFI* and *xoxF* coding sequences reduces the *xoxF* transcripts’ demand for carbon, nitrogen, and hydrogen atoms per codon (Fig. 4). We also show that protein synthesis has been optimized through the frequent incorporation of prebiotic (cheap) amino acids in the pMMO and Xox protein sequences, though not in Mxa (Fig. 3e). Together, these results indicate although the *mxaFI* and *xoxF* coding sequences share similar translation efficiency, the *xoxF* coding sequence reduces transcript and protein synthesis costs (Fig. 5b), suggesting that Xox has evolved to be the predominantly expressed methanol dehydrogenase in environments where oxygen concentration is high and methane concentration is low. Supporting this, recent experimental reports have found that Xox is the preferentially but the *xoxF* coding sequence is specifically expressed when the methane concentration is low^44^. Overall, the results in this study show that the pMMO-Xox configuration of the methane oxidation module should be the most efficient molecular strategy to catalyze the sequential oxidation of methane and methanol (Fig. 5b). This finding complements recent investigations in isolates^43^ and communities^44^ by providing a molecular basis to explain the prevalent expression of *pmoCAB* and *xoxF* coding sequences.

While the genes encoding pMMO have been proposed to have evolved independently in separate genomes following an ancient speciation event^45–47^, it has also been recently suggested that methane oxidation genes may have been acquired by horizontal gene transfer^48,49^. The evolutionary history of the optimized *pmoCAB* coding sequences in type Ia methanotrophs warrants further investigation. Furthermore, hypothesis-driven investigations focusing on the implications of the *pmoCAB* coding sequence in the cellular and environmental performance of methanotrophs will be required. Fundamentally, our results emphasize the type Ia *pmoCAB* coding sequences have been shaped by natural selection and are not the result of random genetic drift even at the level of synonymous codons. The ecological role of methanotrophs as the primary producers in complex communities^5,6^ can exert strong evolutionary pressures on pMMO. Efficient conversion of methane to methanol not only serves as the first enzymatic step for methanotrophs, but also provides methanol as the carbon source for methylotrophic bacteria^32^. The availability of methanol can also support methane oxidation in methanotrophs that have a low affinity for methane when methane concentration is at atmospheric levels^50^. Consequently, the unique adaptations of the type Ia *pmoCAB* coding sequences are likely the result of molecular evolution at the organismal, community, and perhaps even planetary scale^51^.

## Conclusions

By conducting an in-depth meta-analysis of isolate genomes and MAGs from diverse taxonomy and geography we show that, among the seven types of methanotrophs, only type Ia methanotrophs possess a unique coding sequence of the *pmoCAB* operon that is under positive selection for optimal resource allocation and efficient synthesis of transcripts and proteins in environmental counter gradients with high oxygen and low methane concentrations. The coding sequence is conserved in type Ia methanotrophs inhabiting diverse environments around the globe, implying that natural selection rather than neutral evolution governs the molecular evolution of methane oxidation in type Ia methanotrophs. Overall, this study provides genomic evidence of an intriguing adaptive trait in the *pmoCAB* coding sequences of type Ia methanotrophs and extends our knowledge of the evolutionary landscape explored by nature to sustain metabolism in highly diverse environments on Earth.

## Materials and Methods

Detailed description of the methods used in this study is available in Additional file 1: Supplemental Methods.

### Methanotroph genomes

Of the 67 methanotroph genomes analyzed in this study, 49 are isolates and 18 are metagenome-assembled genomes (MAGs) (see Additional file 2 for details). This collection included most of the isolate genomes and relevant methanotrophic metagenomic datasets available in the public domain as of October 2017. Genome annotations were compiled expressed methanol dehydrogenase in isolates of type Ia methanotrophs^43^, while in complex microbial communities, the *mxaFI* from the JGI/IMG^52^ and NCBI/RefSeq^53^ databases. Functional coding sequences of type Ia methanotrophs are preferentially expressed when methane and oxygen are in high concentrations, annotations were absent or ambiguous for the genomes of the type Ia methanotrophs *Methylobacter tundripaludum* SV96, *Methylococcaceae bacterium* Sn10-6, and *Methylomicrobium alcaliphilum* 20Z; type Ib *Methylococcus capsulatus* Bath and *Methylococcus capsulatus* Texas; type IIa *Methylocystis parvus* OBBP and *Methylocystis sp*. SC2; and type III *Methylacidiphilum infernorum* V4. The amino acid sequences from their translated coding sequences were further annotated with KEGG/`BlastKOALA`^54^. Phylogeny of the genomes was assigned by `PhyloPhlAn`^55^ based on the whole genome.

### Metagenomic analysis pipeline

Briefly, raw reads of publicly available metagenomic datasets were obtained from NCBI. Quality control of raw reads was performed with `illlumina-utils`^56^. Contigs were co-assembled with `MEGAHIT`^57^ and binning was performed with `MaxBin`^58^. The quality of MAGs was assessed with `CheckM`^59^ and MAGs were categorized according to the quality standards defined in the Minimum Information about a Metagenome-Assembled Genome of bacteria^60^. Annotation of MAGs was conducted with `Prokka`^61^ and `eggNOG-mapper`^62^.

### Sequence analysis

Nucleotide composition bias (GC and GC_3_ content) were analyzed with `seqinR`^63^ in R. Codon usage metrics were formulated in R according to the equations described for RSCU^64^ and CAI^64^. tAI was calculated with `codonR`^65^. The tRNA datasets required for reproduction of tAI calculations can be found in Additional file 1: Figure S10. ENC^66^ was calculated with chips in `EMBOSS`^67^.

### Network analysis

The network dataset was generated using in-house scripts. The network matrix was imported and edited with `Cytoscape`^68^.

### Transcriptomic analysis

Normalized mRNA abundance data from RNA-Seq experiments were obtained from three publicly available independent studies conducted in the type Ia methanotrophs *Methylomicrobium buryatense* 5G^31^, *Methylomicrobium alcaliphilum* 20Z^30^ and *Methylobacter tundripaludum* 31/32^32^.

### Code and data availability

Scripts required to reproduce all the results and figures can be obtained from https://github.com/PLeeLab/methane_oxidation_genetic_trait/.

## Declarations

## Acknowledgements

This research was supported by the Research Grants Council of Hong Kong through Project 11206514. JCV acknowledges support provided by the Hong Kong PhD Fellowship Scheme (HKPFS). We thank David Wilkins for critiquing the manuscript.

