## Additional file 1 for "The unique coding sequence of *pmoCAB* operon from type Ia methanotrophs simultaneously optimizes transcription and translation"

### Supplemental Methods

**Computational packages and scripts.** *R* [1], *RStudio* [2], and *ggplot2* [3] were used to produce all the analyses and figures presented in this study unless otherwise indicated. All the scripts used in this work are available at GitHub (https://github.com/PLeeLab/methane_oxidation_genetic_trait).

**Metagenomics analysis pipeline**. Our analysis was applied to publicly available metagenomic data from five potentially methanotrophic environments: 1) Lake Washington, USA [4]; 2) Serpentinite Springs of the Voltri Massif, Italy [5]; 3) Movile Cave in Mangalia, Romania [6]; 4) Santa Elena Ophiolite alkaline spring, Costa Rica [7]; and 5) Coastal basin of Golfo Dulce, Costa Rica [8]. For 1), the publicly available metagenome-assemble genomes (MAGs) (https://gold.jgi.doe.gov/studies?id=Gs0114290) were also examined.

The metagenomics analysis pipeline consists of seven main stages and a preliminary Stage 0 for data and software preparation. In Stage 0, raw metagenomic reads in FASTQ format were downloaded from NCBI/SRA using *fastq-dump* from the SRA Toolkit [9]. When a sample was produced using paired-end sequencing, sample integrity was verified by confirming it contained the same number of forward and reverse reads. The average and standard deviation read count were then calculated. The algorithms and packages used in our pipeline are summarized below:

| Stage | Function | Software | Ref | Link |
|---|---|---|---|---|
| 0 | Preliminary | *fastq-dump* | [9] | https://ncbi.github.io/sra-tools/fastq-dump.html |
| 1 | Quality control | *illlumina-utils* | [10] | https://github.com/merenlab/illumina-utils |
| 2 | Co-assembly | *MEGAHIT*<br>*Anvi'o* | [11]<br>[12] | https://github.com/voutcn/megahit<br>https://github.com/merenlab/anvio |
| 3 | Binning | *MaxBin* | [13] | https://downloads.jbei.org/data/microbial_communities/MaxBin/MaxBin.html |
| 4 | Refine bins | *CheckM* | [14] | http://ecogenomics.github.io/CheckM/ |
| 5 | Functional annotation | *Prokka* | [15] | https://github.com/tseemann/prokka |
| 6 | Taxonomy classification of bins | *PhyloPhlAn* | [16] | https://bitbucket.org/nsegata/phylophlan/wiki/Home |
| 7 | Refine functional annotation of | *eggNOG-mapper* | [17] | https://github.com/jhcepas/eggnog-mapper |

| | |
|---|---|
| | methanotrophic MAGs |

For Stage 1 (quality control), high-quality reads were selected using *illlumina-utils* [10] with the Minoche [18] method using the command *iu-filter-quality-minoche* with the parameter *--ignore-deflines*. As the only exception, we used the method *iu-merge-pairs* for the metagenome from Serpentinite Springs of the Voltri Massif, Italy as the authors reported that the sequencing in this project yielded partially overlapped paired-end reads. In Stage 2 (co-assembly), pooled samples from the same environment were co-assembled following published methods [19–21]. Briefly, reads that passed quality control were co-assembled into contigs with *MEGAHIT* using the parameter *--min-contig-len* = 1000. Contigs produced by *MEGAHIT* [11] were subjected to refinement with *anvi'o* [12] three times using the parameter *--simplify-names* and setting *--min-len* to 1000, 1500, and 2500. In Stage 3 (binning), refined contigs and high-quality reads were binned with *MaxBin* [13] to produce MAGs (MAGs and bins refer to the same item in this pipeline). In Stage 4 (refine bins), the quality of MAGs was assessed with *CheckM* [14] and retained if they exceeded the quality standards defined in the Minimum Information about a Metagenome-Assembled Genome (MIMAG) for bacteria [22] for a medium-quality draft (completeness > 70% and contamination < 10%). The exception was the methanotroph MAG from the Movile Cave, Romania [6] which had a completeness of 35%. We included this MAG for comprehensiveness of MAGs assembled from environmental sources. MAGs of potential methanotrophs were selected according to the presence of methane oxidation genes (*pmoCAB* or *mmoXYZCDB*) in their genomes. In Stage 5 (functional annotation), MAGs were annotated with *Prokka* [15] using the parameters *–metagenome* and *--kingdom=Bacteria*. In Stage 6 (taxonomic classification), the amino acid sequences produced by *Prokka* were used as input for taxonomic characterization of MAGs using *PhyloPhlAn* [16] with parameters *-i* and *-t*. Only MAGs resulting in a taxonomic classification with high or medium confidence were selected for subsequent analyses. Finally, in Stage 7 (functional annotation refinement), all MAGs characterized as probable methanotrophic bacteria were subjected to a second, more comprehensive annotation procedures with *eggNOG-*

*mapper* [17], in which KEGG Orthologs (KO), Gene Ontology (GO), and Clusters of Orthologous Groups (COGs) were assigned to genome features.

The retrieved publicly available assembled and binned MAGs of methanotrophic bacteria of Lake Washington, USA [4] were subjected to our metagenomics analysis pipeline from Stage 4 to Stage 7.

**Geographical location of methanotroph genomes**. The geographical coordinates of the origin for each sample were determined either from manual inspection of published reports (Additional file 2) or IMG/JGI [23]. When available, the exact coordinates of the sampling location were used to place genomes in the map. For nine genomes, the origin location could not be found using either method. The coordinates were plotted using the *maps* [24] package in *R*. The *position_jitter* parameters were set to $w = 3.1$and $h = 3.1$ to avoid overlapping of dots.

**Genome-scale phylogenetic tree of genomes and MAGs.** 67 methanotroph genomes and MAGs and one outgroup genome of the non-methanotrophic bacterium *Bacteroides ovatus* ATCC 8483 were used to reconstruct the phylogenetic tree with *PhyloPhlAn* [16] with parameter *-u* (*de novo* phylogenetic tree).

**Incorporating metadata and nucleotide content into genome-scale phylogeny.** The resultant phylogenetic tree (raw tree *1_proteomes_tree.nwk* available in GitHub repository) was imported to *R* using the *ape* package [25] and re-rooted to the outgroup genome of *B. ovatus* ATCC 8483. The outgroup genome was selected based on its close placement to known methanotrophs in the microbial tree of life [26]. Metadata of genomes and MAGs were also imported in order to assign features to each genome and to differentiate the seven methanotroph types. Methanotroph types were assigned using the *treeio* [27] *R* package. The tree, number of coding sequences (CDSs) and distribution of GC and $GC_3$ content were plotted using the *R* packages *ggtree* [28] and *ggridges* [29]. GC and $GC_3$ compositions of CDSs were determined using the *gc* and *gc3* functions of the *seqinr* [30] *R* package. The standalone version of *EMBOSS* [31] was used to corroborate the GC and $GC_3$

content of each CDS of our interest. All the data were compiled and manually curated and are available in the file *1_QC_CH4.txt* in our GitHub repository.

**Analysis of relative synonymous codon usage (RSCU).** The frequency of individual codon usage per CDS normalized to the amino acid usage of its corresponding protein was calculated as RSCU [32] using the function *uco* with parameter *index = rscu* from the *seqinr R* package. The equation used to calculate the RSCU is:

$$RSCU = \frac{O_{ij}}{\frac{\left[\sum_{j}^{n_i} O_{ij}\right] * 1}{n_i}} \quad (1)$$

where $O_{ij}$ is the occurrence of the *j*th codon for the *i*th amino acid and $n_i$ the total number of synonymous codons coding for the *i*th amino acid. We considered a codon frequently used if RSCU ≥ 1.6, and rarely used if RSCU ≤ 0.6. Principal component analysis (PCA) was computed using the *R* function *prcomp* to identify CDSs that share similar preferences codon usage biases based on RSCU values for 59 codons (the conventional set of 64 codons excluding the two non-redundant codons for methionine and tryptophan, which have a fixed RSCU = 1.0, and the three stop codons).

**Calculation of the codon adaptation index (CAI)**. The CAI [32] was used to analyze the codon usage of each CDS relative to a reference set of CDSs. Codon frequency was calculated for each CDS in each of the 67 isolate genomes and MAGs. Frequencies were calculated for a single reading frame of the CDS and only ~1% of all CDSs had length not divisible by three. The codon relative adaptiveness (*w*) was calculated as the frequency of a codon divided by the frequency of the synonymous codon with the highest frequency [32]. *w* values were used to compute the CAI for each codon using the *cai* function from the *seqinr R* package, using either the full set of CDSs in the genome ($CAI_{genome}$) or only ribosomal protein genes

($CAI_{ribosome}$). The percentile rank of each CDS within the distribution of $CAI_{genome}/CAI_{ribosome}$ was calculated.

**Analysis of the effective number of codons (ENC).** ENC [33] is a measure of CDS codon usage bias based on codon preference per amino acid and has been applied recently to study genomes assembled from environmental samples [34, 35]. ENC values were computed for each CDS using the *chips* program from *EMBOSS* [31] based on Wright's equation [33]:

$$ENC = 2 + \frac{9}{\hat{F}_2} + \frac{1}{\hat{F}_3} + \frac{5}{\hat{F}_4} + \frac{3}{\hat{F}_6} \tag{2}$$

where $\hat{F}_i$ is the codon homozygosity for the amino acids of degeneracy *i*. ENC as a function of $GC_3$ content was analyzed in all methanotrophs. A linear model relating ENC and $GC_3$ was fitted using the *stat_smooth* function from the *ggplot2* R package.

**tRNA copy number.** tRNA frequencies were analyzed only for isolate genomes where the total tRNA pool should be known. When available, the tRNA counts of the genomes were downloaded from the public databases of IMG/JGI and GtRNAdb [36, 37], otherwise they were computed with the local version of tRNAscan-SE 2.0 [38].

**tRNA adaptation index (tAI).** tAI was developed to estimate translation efficiency [39, 40]. The tAI was calculated for all CDSs in each genome using the *R* package *codonR* [40] with the parameter *sking* set to *1* (Prokaryote super kingdom) and the default *s* parameter for codon selection penalties. The tAI computation required the genomic tRNA counts (Additional file 1: Figure S10) and CDS codon frequencies, which were calculated using *CodonM*. Within-genome tAI percentile ranks were calculated for each CDS. The manually curated dataset containing the tAI data for our CDSs of interest can be found in the file *1_QC_V_manuallycurated.txt* in our GitHub repository.

**Interaction network of codons and tRNAs.** The interaction network was reconstructed for isolates of type Ia methanotrophs based on RSCU values for six CDS sets, tRNA copy numbers and codon-anticodon pairing rules. Four CDS sets (*pmoCAB*, *mmoXYZCDB*, *mxaFI* and *xoxF*) represented the methane oxidation metabolic module, one set comprised ribosomal protein genes, and one set comprised all the CDSs in each genome. The median RSCU of each codon for each CDS set was computed from the distribution of RSCU values of all type Ia methanotrophs. The median copy number of each tRNA anticodon was calculated from all the copy numbers of all tRNA anticodons in type Ia methanotrophs. The tRNA anticodon matrix is shown in Additional file 1: Figure S10. Standard codon-anticodon recognition rules [40] were used and are detailed in *0_wobble_pairing_rules.txt* available in our GitHub repository.

The integrated dataset was transformed into a network of sources (tRNA anticodons) and targets (CDS codons). The raw network matrix can be found in the file *RSCU_complete_network.txt* in GitHub. The matrix was imported to Cytoscape [41] and edited as shown in the Cytoscape file *1_Fig3D_net_cytoscape.cys*. The complete network containing all amino acids and codons is shown in Additional file 1: Figure S11. A quantitative analysis was applied to the raw network matrix (*RSCU_complete_network.txt*). For each CDS, the number of accessible tRNA copies was calculated for a range of RSCU thresholds (e.g. for RSCU threshold = 0 each CDS can access every possible tRNA). This allowed the number of tRNA copies that a CDS can access as function of codon bias usage (as determined by RSCU) to be calculated. For each CDS, access to the tRNA pool can be measured in absolute term and relative to the tRNA pool available when compared with the access granted to other CDSs. The significance of the difference between two states (RSCU = 0 and RSCU = 2) was assessed with a Chi-square test, with accessible tRNA copies at RSCU = 0.0 as the expected value and at RSCU = 2.0 as the observed value. The test was applied only to CDSs of the methane oxidation metabolic module.

**CDS amino acid composition.** For each CDS, the codon exhibiting the highest median RSCU for each amino acid was selected. Methionine and tryptophan were excluded as they are each encoded by only one codon. The median and standard deviation were calculated from the distribution of RSCU values of each type of methanotrophs. The median and standard deviation of amino acid composition of each translated CDS of each operon was calculated using the distribution of the amino acid composition of each type of methanotrophs. A linear model (using the *lm* function in *R*) was fitted to determine the relationship between codon preference and amino acid usage to serve as a proxy to identify selection for optimal codons at synonymous sites occupied by the most abundant amino acids.

**Prebiotic amino acids analysis.** The amino acid content of each protein in the metabolic module of all methanotrophs was calculated from its translated CDS. Amino acids were categorized as ‘cheap’/prebiotic (alanine, aspartic acid, glutamic acid, glycine, isoleucine, leucine, proline, serine, threonine, and valine) or ‘expensive’/modern amino acids [42–44]. A *t*-test was used to compute the statistical significance of the difference between modern and prebiotic amino acid composition of each protein in the metabolic module from type Ia methanotrophs. The sample for this test was the median amino acid composition (Additional file 1: Figure S12).

**Transcriptome analysis**. Transcribed CDSs were analyzed by modifying thymine (T) for uracil (U) in all CDSs. Three publicly available type Ia methanotroph transcriptomic datasets (*Methylomicrobium buryatense* 5G [45], *Methylomicrobium alcaliphilum* 20Z [46] and *Methylobacter tundripaludum* 31/32 [47]) were used. Normalized mRNA abundance was obtained from each dataset as reported. The purine (A+G) and pyrimidine (T+C) content of each transcribed CDS was calculated. The purine and pyrimidine content of each transcriptome was calculated based on the ribonucleotide composition of each transcribed CDS multiplied by the transcript abundance, summed across all transcribed CDSs. The effect on transcriptome composition of removing a set of transcripts was calculated by subtracting the total transcribed CDS composition (transcribed CDS composition × transcript abundance) from the dataset and re-calculating the total ribonucleotide composition.

**Elemental composition of transcribed CDSs.** The carbon (C), hydrogen (H), oxygen (O) and nitrogen (N) composition of transcribed CDSs was calculated based on ribonucleotide molecular formulae (adenine $C_5H_5N_5$, guanine $C_5H_5N_5O$, cytosine $C_4H_5N_3O$, uracil $C_4H_4N_2O_2$) and normalized to the number of codons in each CDS. To provide statistical support for the observations of elemental composition bias in *pmoCAB* transcripts, the mean per-codon elemental content of 1,000 randomly selected combinations of three transcribed CDSs was calculated.

**Correlation between transcript abundance and elemental composition.** It has been recently proposed that highly expressed genes tend to decrease per-codon nitrogen requirements of their RNA transcripts [48, 49]. The relationship between elemental composition and mRNA abundance was investigated by computing the Pearson correlation coefficient with 95% confidence levels.

#### Supplemental Figures

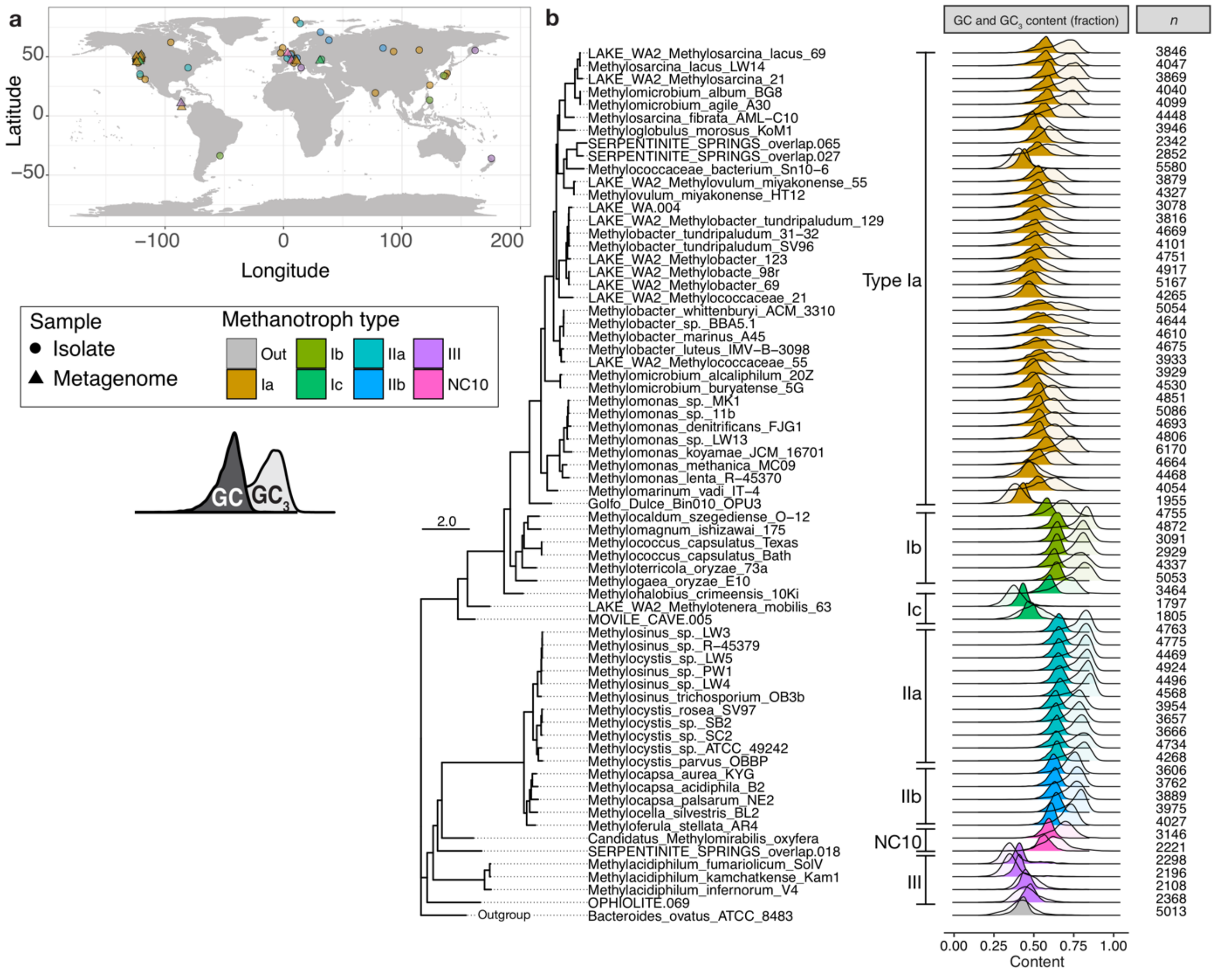

**Figure S1.** Nucleotide composition bias of genomes. (a) Genomes analyzed in this study were geographically mapped to the locations where the samples originated. (b) Phylogenetic tree was reconstructed based on whole-genome sequences. The total number of coding sequences (*n*) used to calculate the statistics of GC and $GC_3$ content for each organism is indicated.

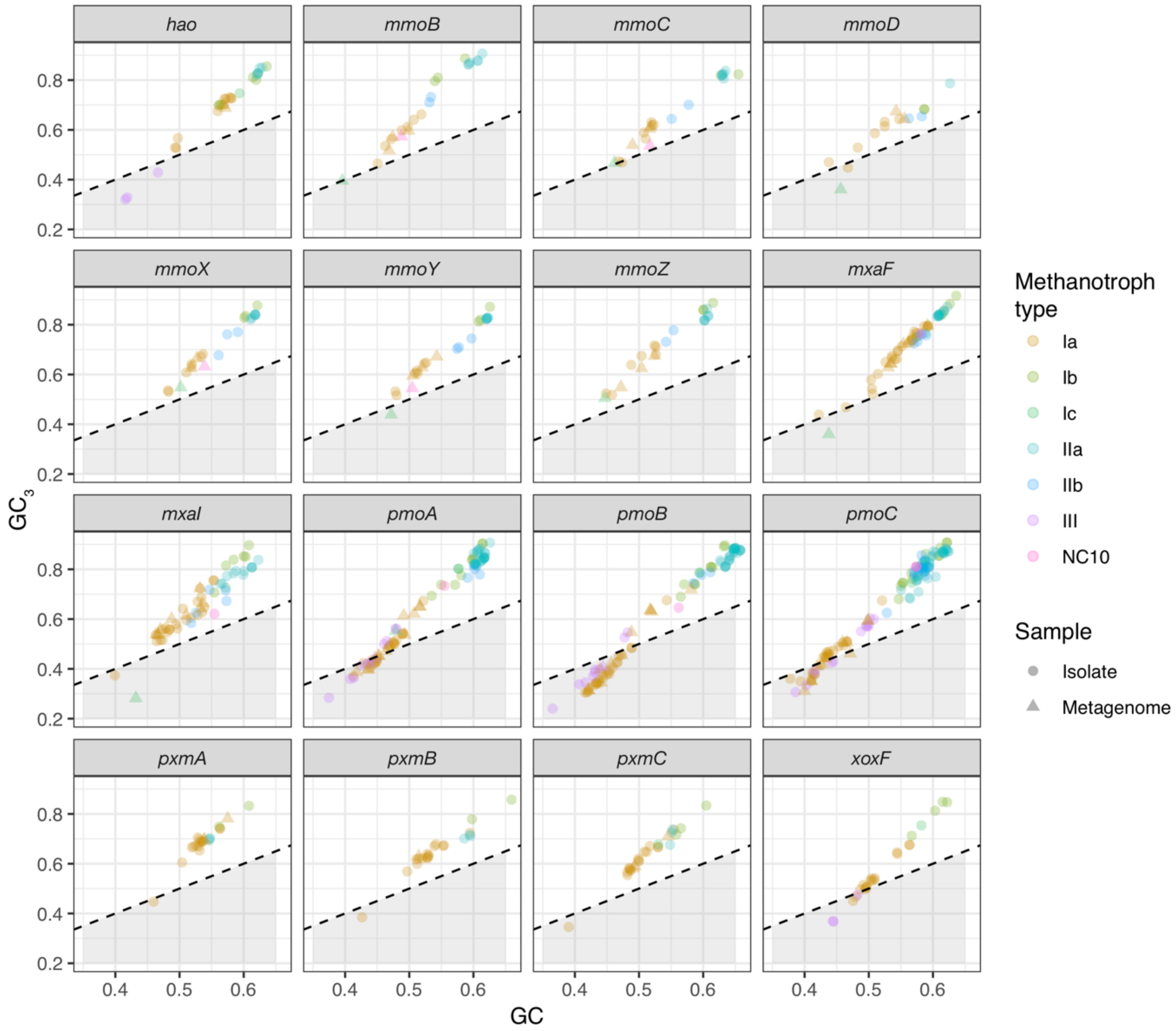

**Figure S2.** An extended analysis of the nucleotide composition bias of all CDSs in the methane oxidation metabolic module and controls (*hao* and *pxmABC*) in all methanotrophs. The dashed line indicates equality between GC and $GC_3$. The shaded region below the equality line shows the data points where $GC_3 < GC$.

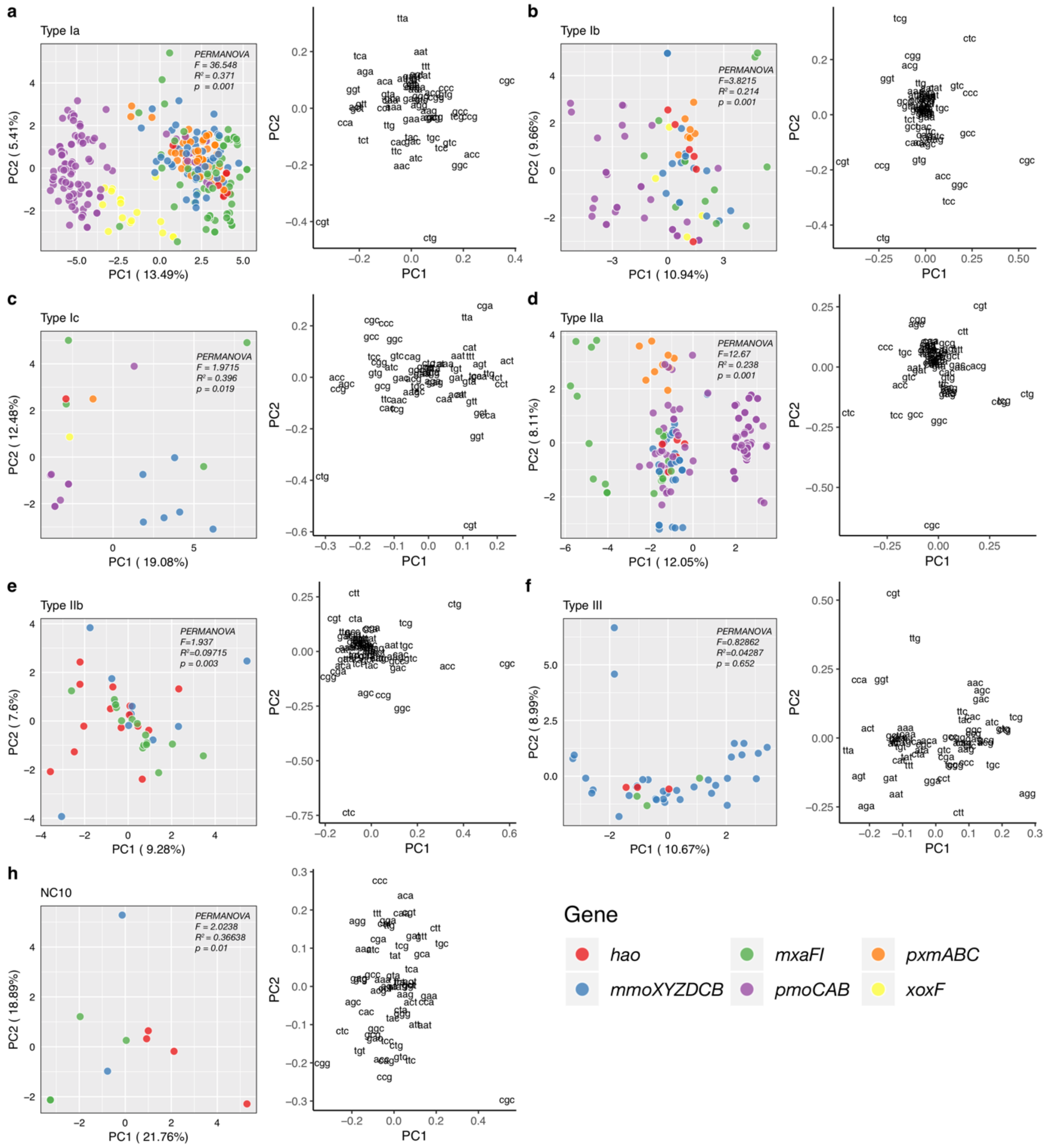

**Figure S3.** An extended analysis of the RSCU bias of all CDSs in the methane oxidation metabolic module and controls (*hao* and *pxmABC*) in all methanotrophs. Principal component analysis (PCA) of the RSCU values for the four CDSs of the methane oxidation metabolic module and two controls of (a) type Ia, (b) type Ib, (c) type Ic, (d) type IIa, (e) type IIb, (f) type III, and (g) NC10 methanotrophs. Plots at the right-hand side of each PCA show the variable (codons) loadings of the PCA.

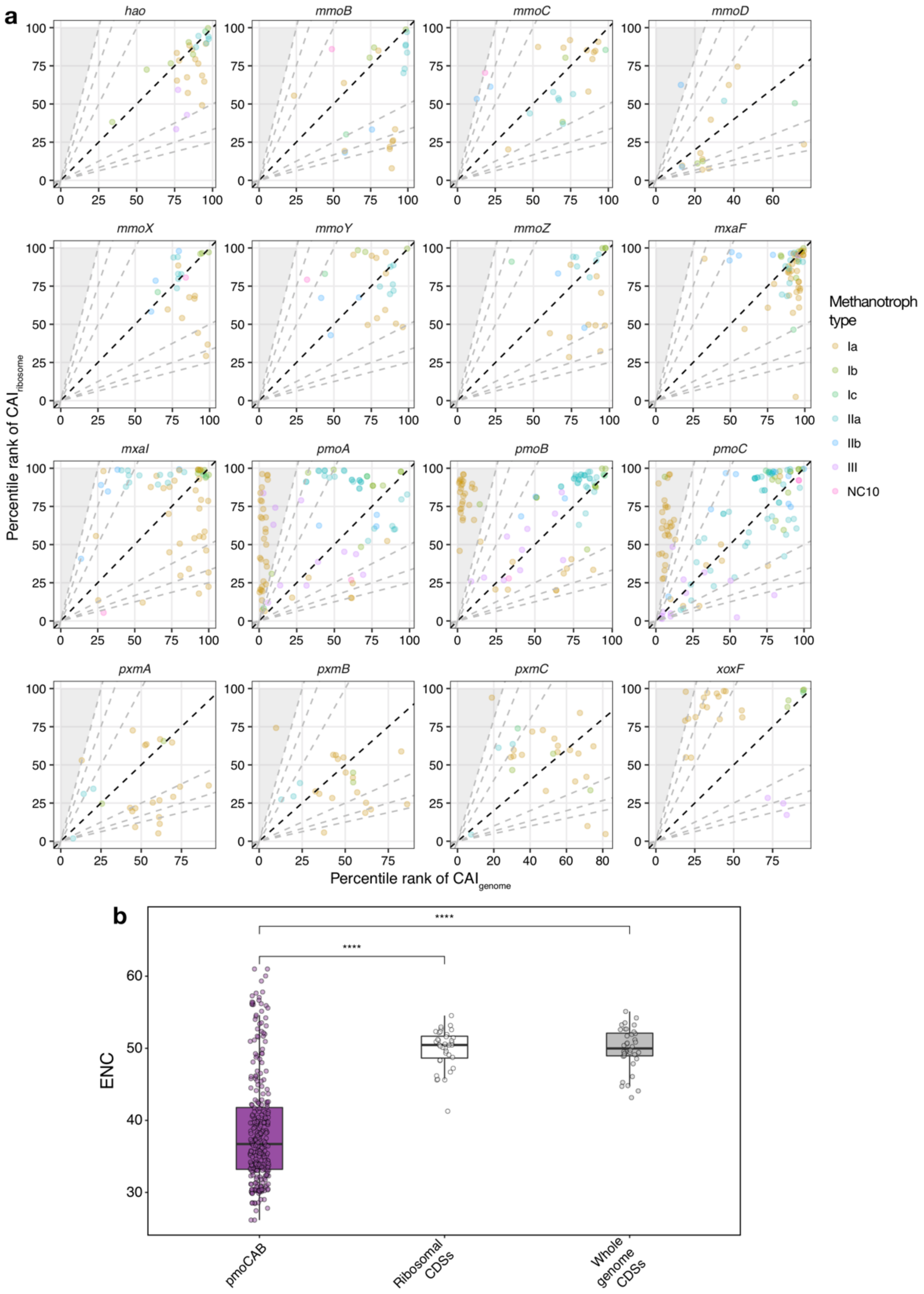

**Figure S4.** An extended analysis of CAI and ENC. (a) CDSs in the methane oxidation metabolic module and controls (*hao* and *pxmABC*) in all methanotrophs. CAI indexes were converted to percentile ranks based on the relative distribution of CAI in each methanotroph. Dashed lines with varying slopes delineate the variations between the percentile ranks of both indexes. (b) Comparison between the ENC of CDSs of *pmoCAB*, ribosomal proteins, and whole genomes of type Ia methanotrophs. The two-tailed Wilcoxon signed rank-test with 99% confidence level was used to test the significance of the difference. ****$p \leq 0.0001$.

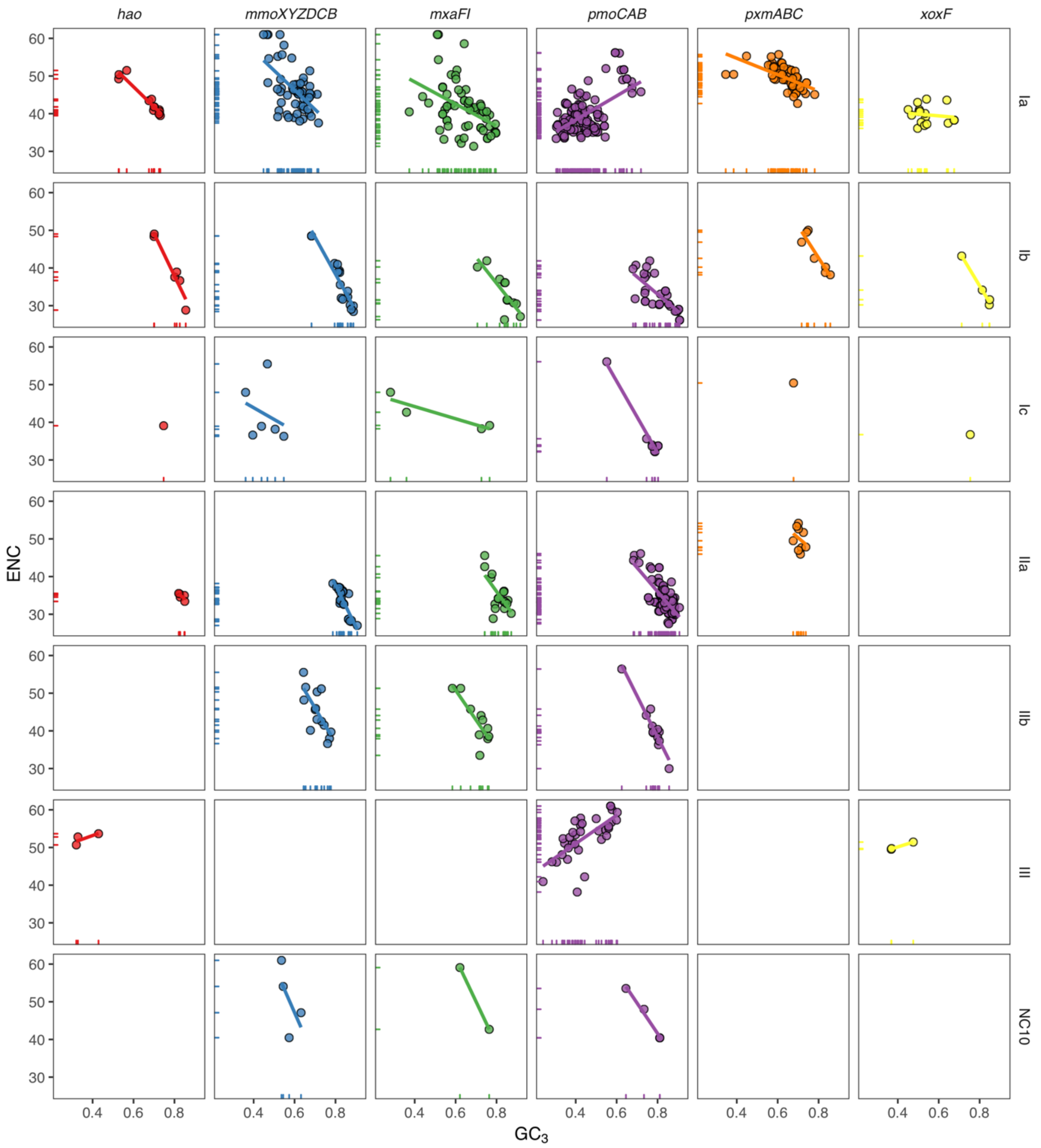

**Figure S5.** An extended analysis of the ENC as a function of $GC_3$ of all CDSs in the methane oxidation metabolic module and controls (*hao* and *pxmABC*) in all methanotrophs. The line represents the fit of a linear model with formula ENC ~ $GC_3$.

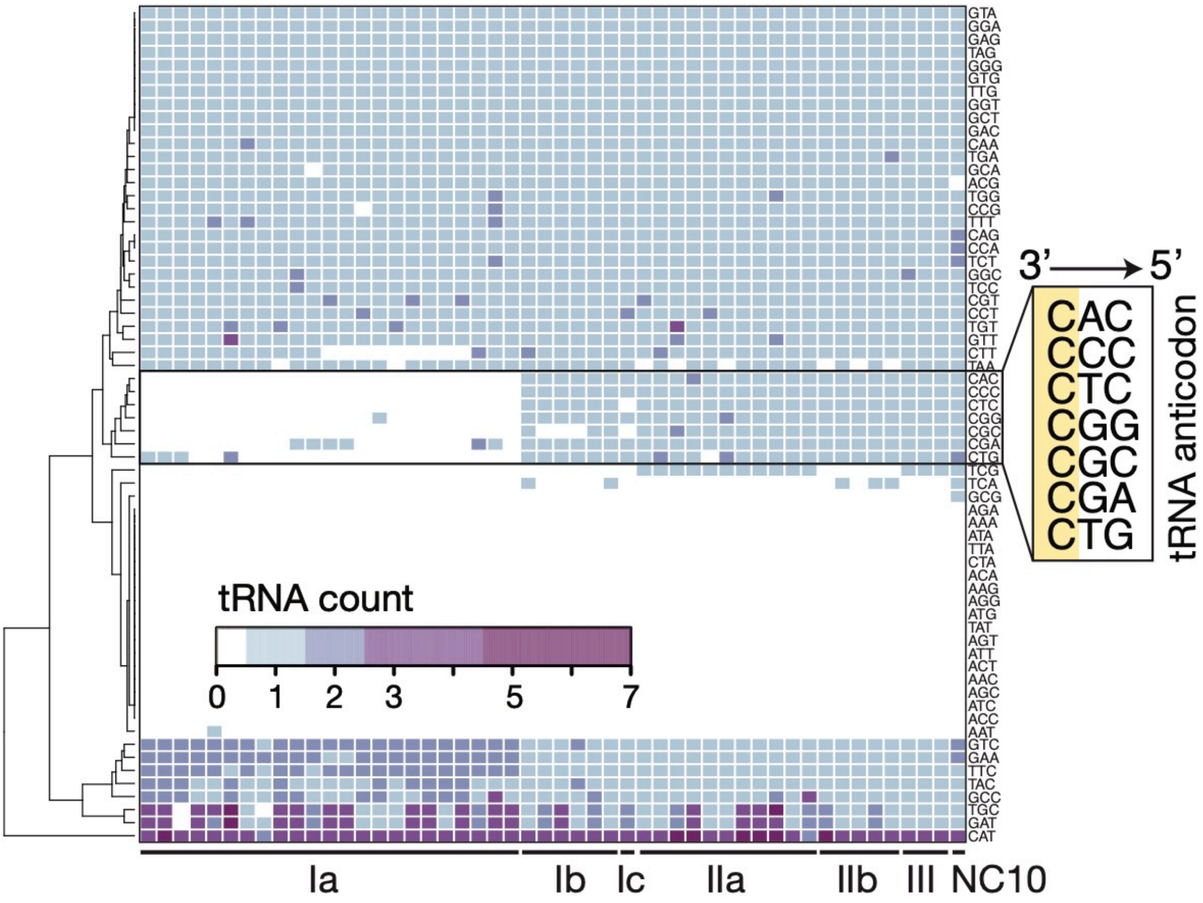

**Figure S6.** The tRNA pool of each genome.

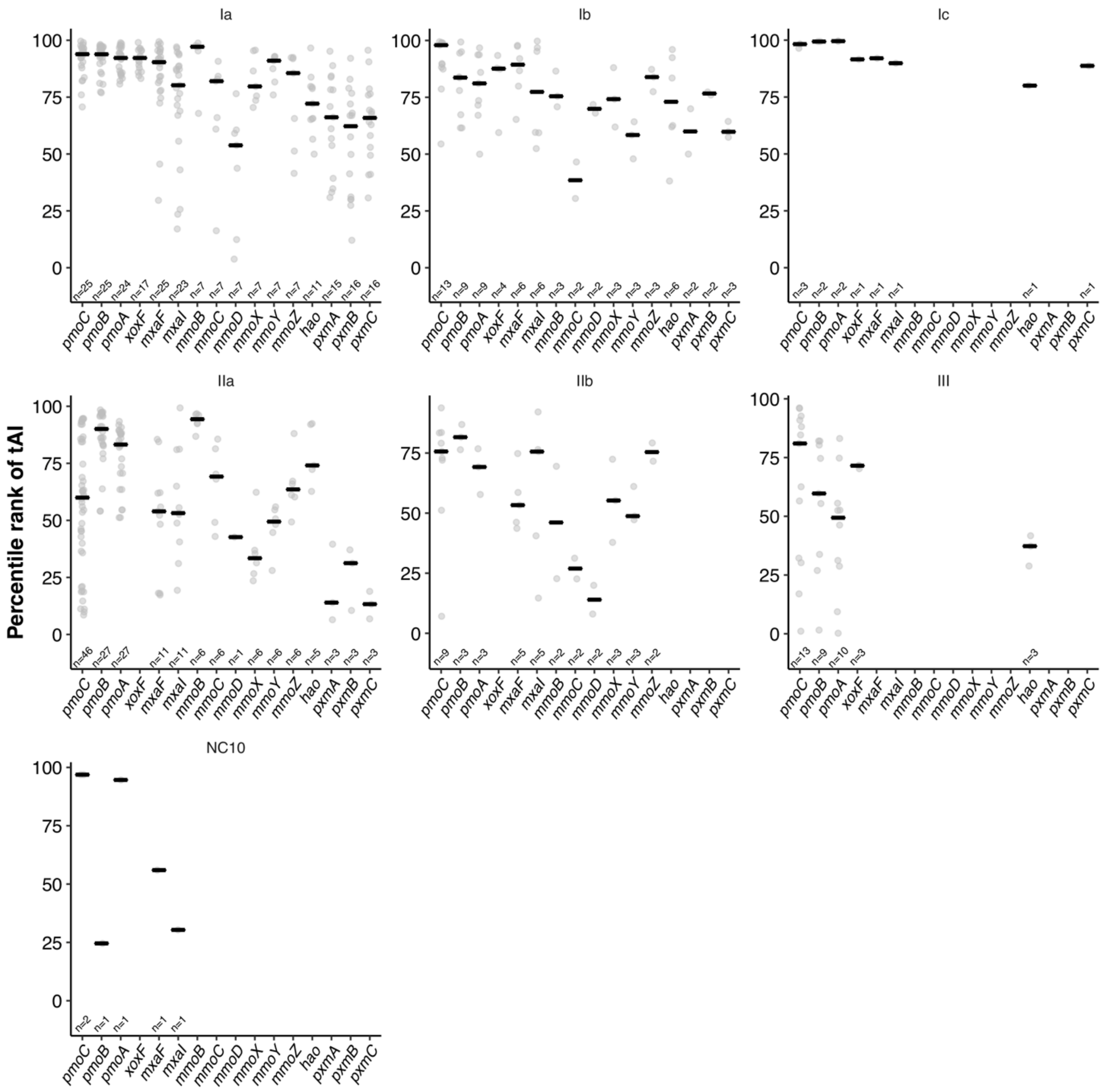

**Figure S7.** An extended analysis of the tAI of all CDSs in the methane oxidation metabolic module and controls (*hao* and *pxmABC*) in all methanotrophs. tAI values were converted to percentile ranks based on the relative distribution of tAI in each methanotroph. The black horizontal line represents the median tAI. *n* is the number of CDSs.

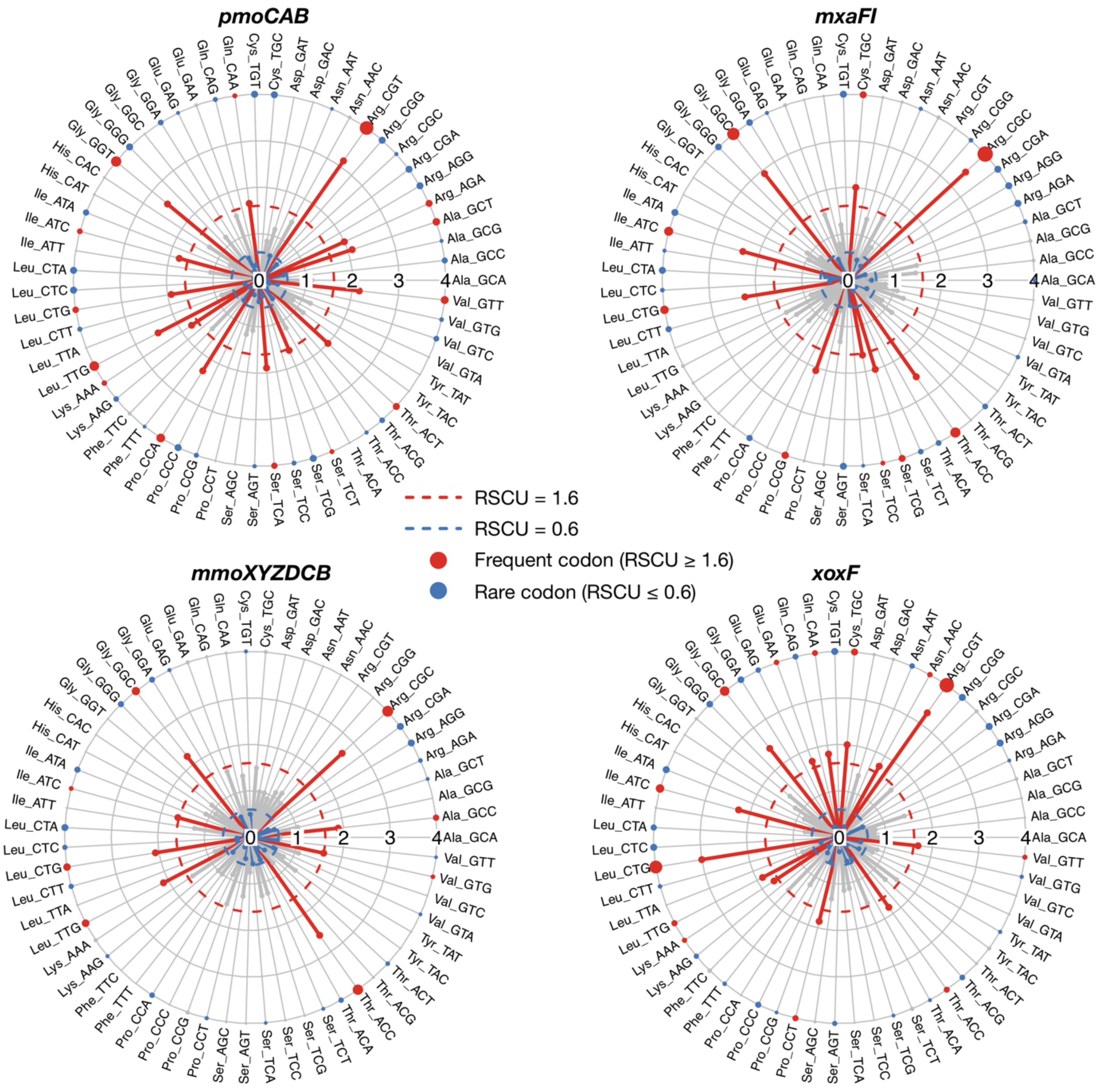

**Figure S8.** Median RSCU values calculated from the distribution of each codon RSCU values in all type Ia methanotrophs.

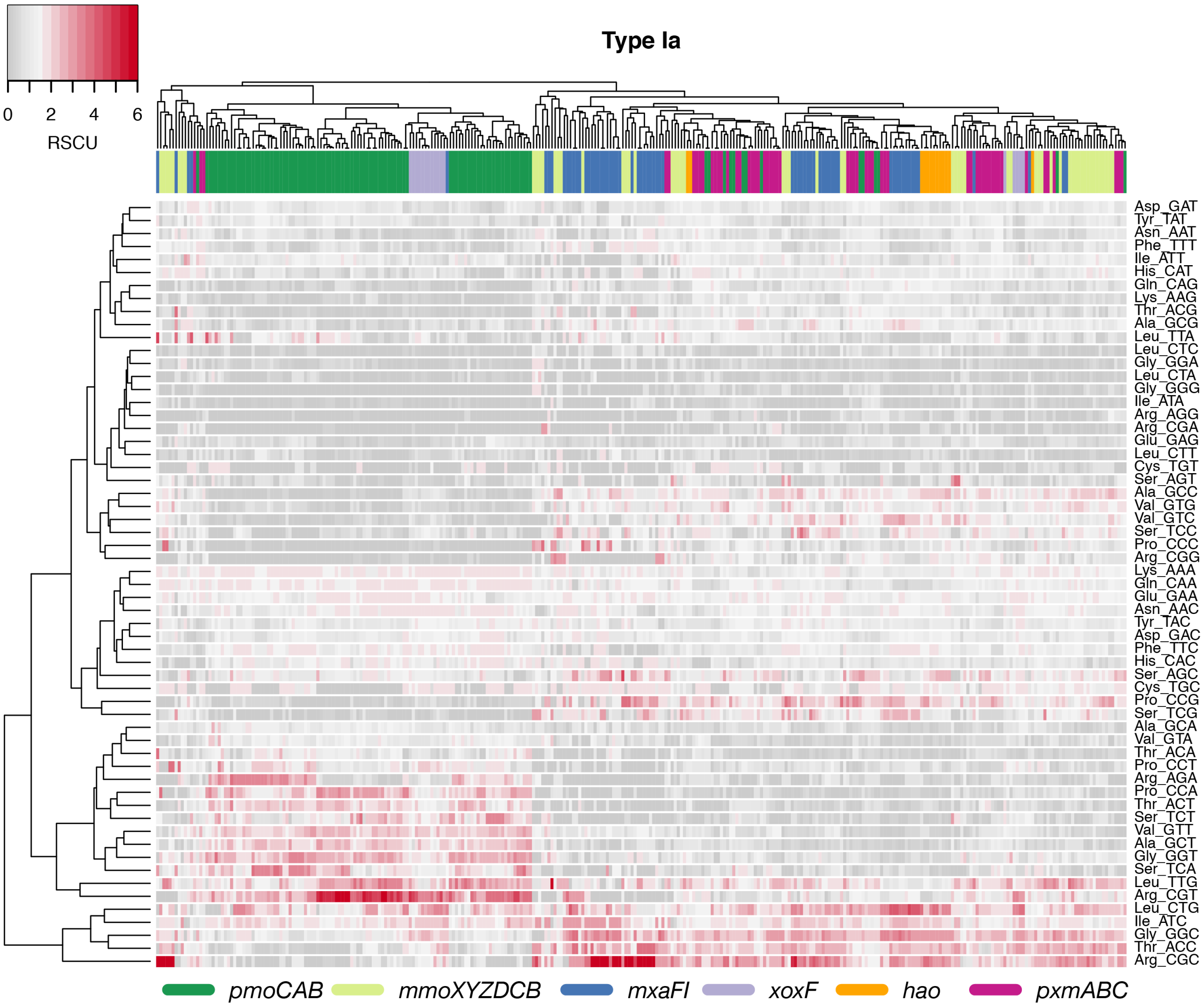

**Figure S9.** A detailed distribution of RSCU values of all CDSs in the methane oxidation metabolic module and controls (*hao* and *pxmABC*) of type Ia methanotrophs. CDSs of each type Ia methanotroph are in the columns and codons in rows.

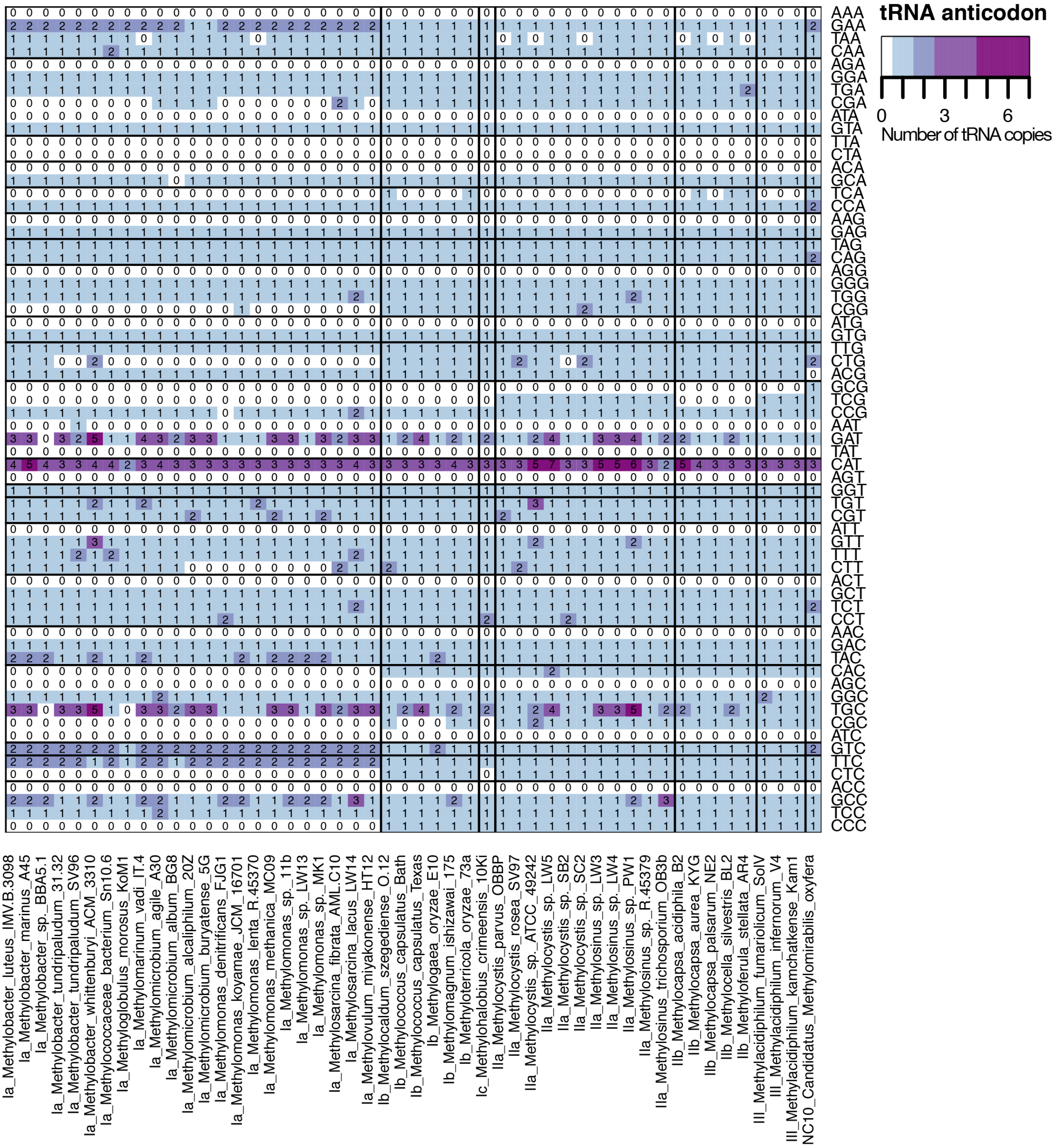

**Figure S10.** The tRNA pool of each genome. Vertical lines divide the different types of methanotrophs. The type of methanotroph is specified in the labels at the bottom.

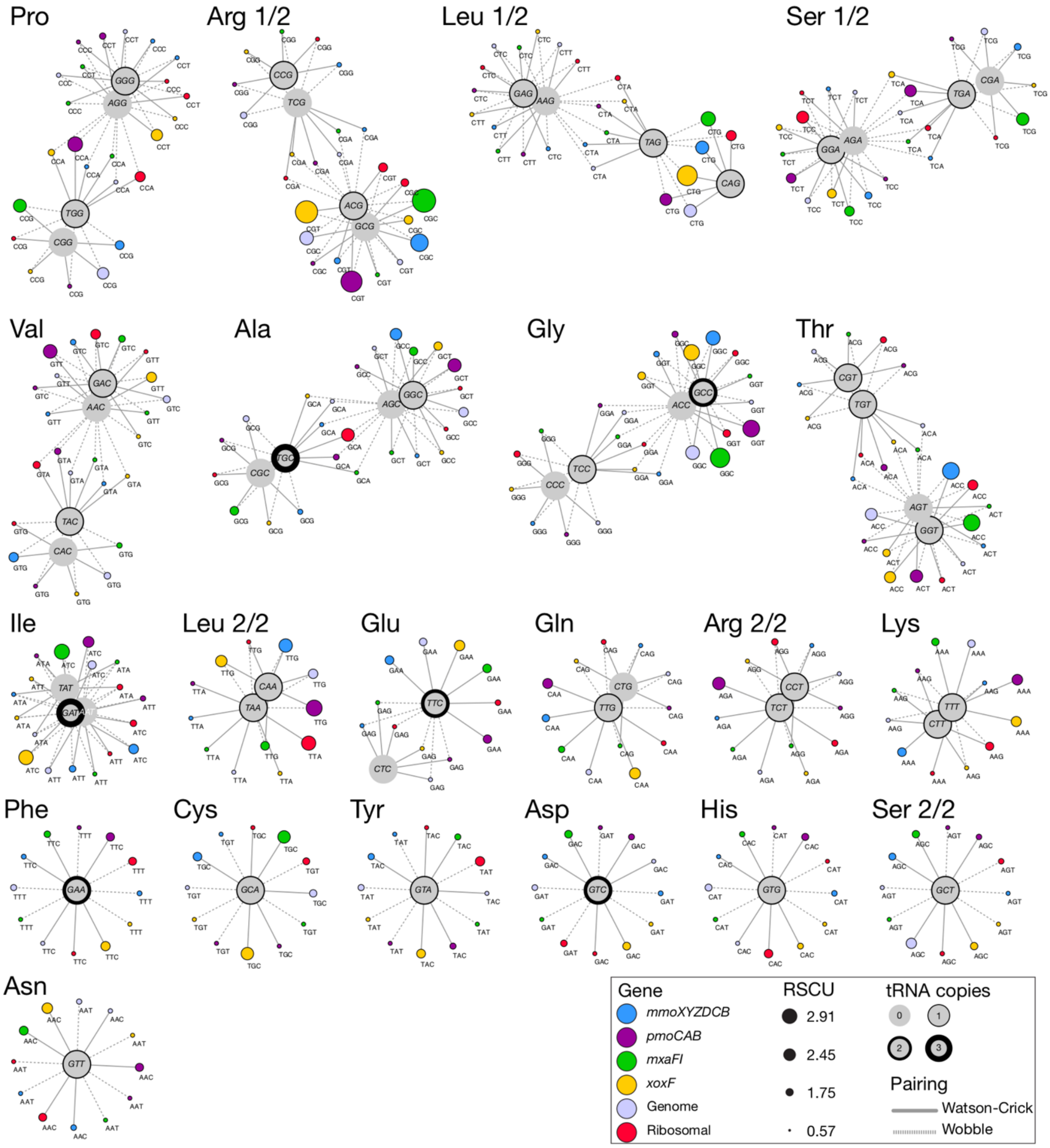

**Figure S11.** An extended network of the interaction between codons and tRNA in type Ia methanotrophs. The integrated network is composed by median RSCU values, median number of tRNA copies, and the codon-anticodon pairing rules. The median of the RSCU values was calculated for six sets of CDSs: four sets in the methane oxidation metabolic module, one set comprising the CDSs of ribosomal protein genes, and one set comprising all the CDSs in the genome.

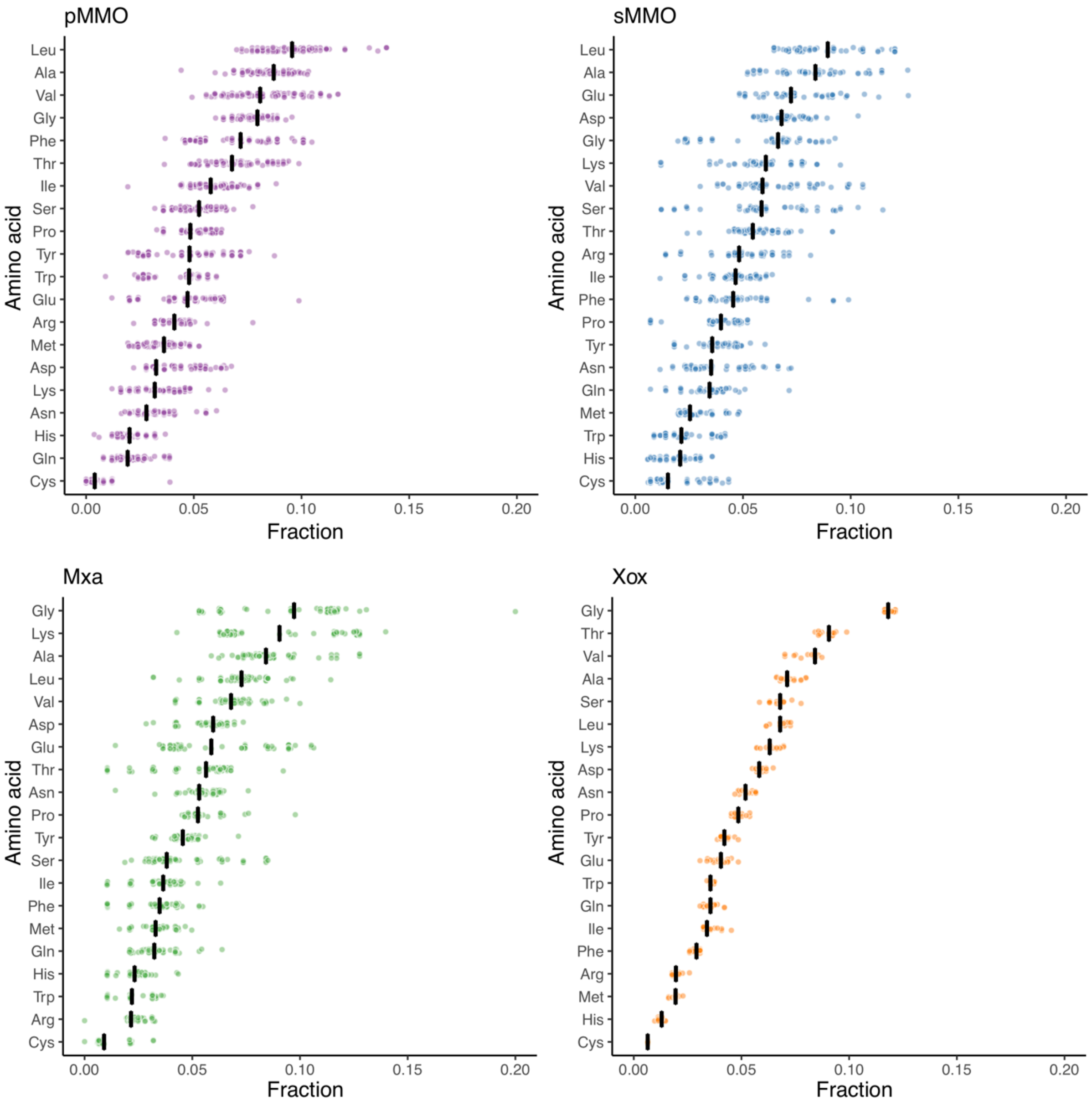

**Figure S12.** Median amino acid usage of each translated CDS of type Ia methanotrophs. The vertical black line in each amino acid shows the median of the amino acid composition as the fraction of the total protein composition.

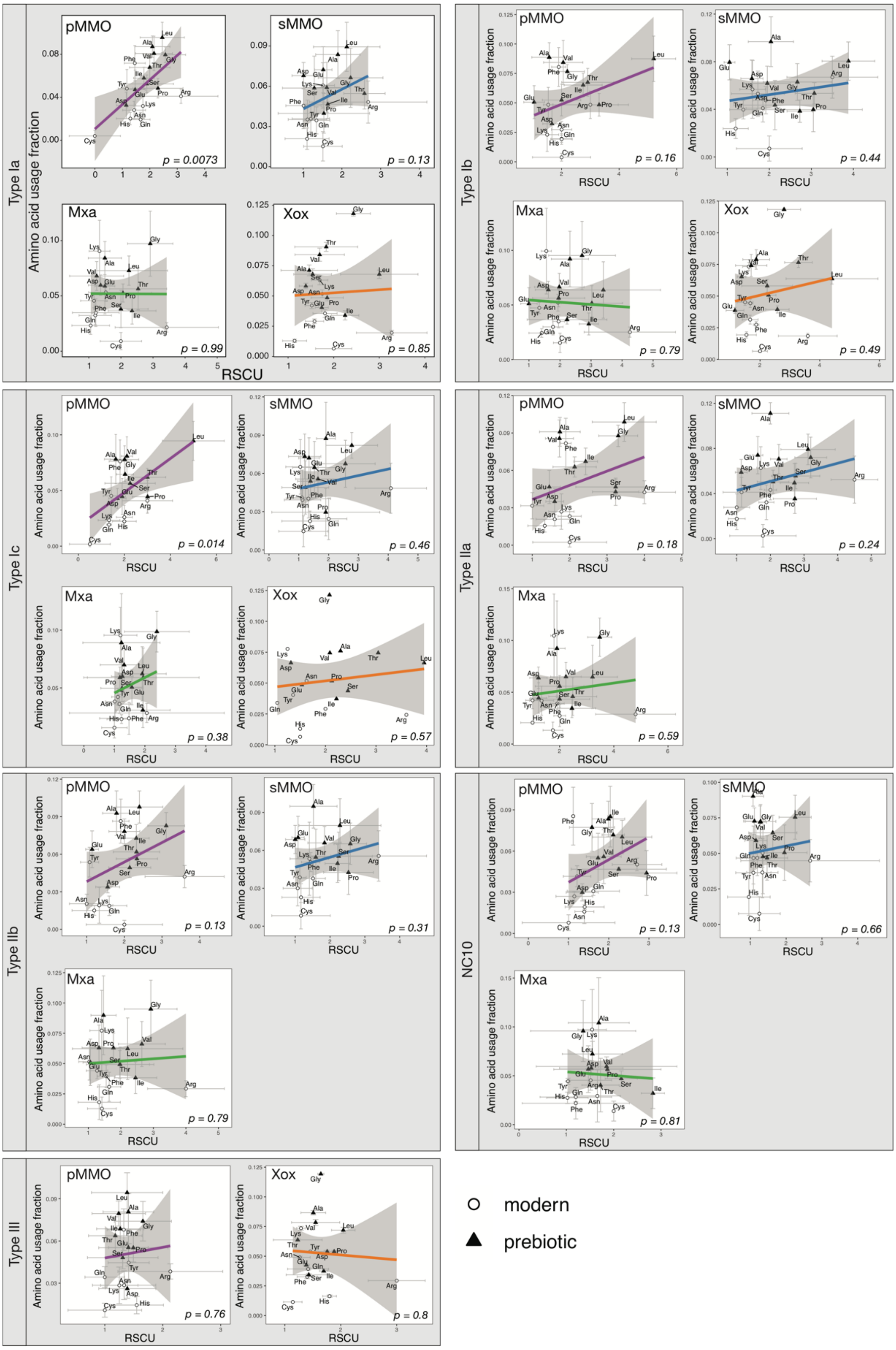

**Figure S13.** Median usage of each amino acid in each translated CDS as a function of the median RSCU value of the codon exhibiting the highest preference for each amino acid for type Ia, Ib, Ic, IIa, IIb, III, and NC10 methanotrophs. The *p*-value corresponds to the significance of the linear regression.

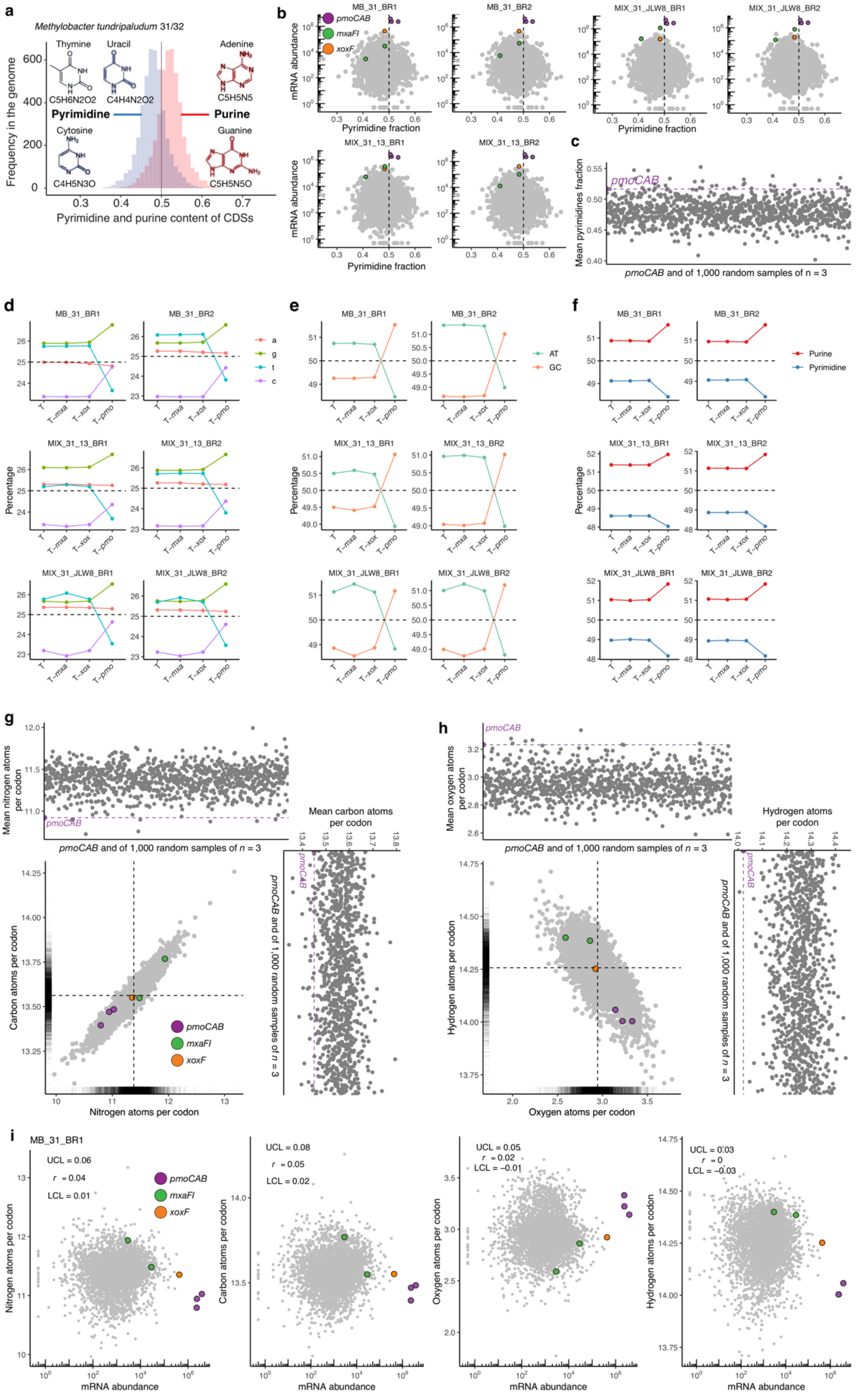
a
Methylobacter tundripaludum 31/32
Thymine
Uracil
Adenine
C5H6N2O2
C4H4N2O2
C5H5N5
Pyrimidine
Purine
Cytosine
Guanine
C4H5N3O
C5H5N5O
Frequency in the genome
Pyrimidine and purine content of CDSs
b
MB_31_BR1
MB_31_BR2
MIX_31_JLW8_BR1
MIX_31_JLW8_BR2
MIX_31_13_BR1
MIX_31_13_BR2
pmoCAB
mxaFI
xoxF
mRNA abundance
Pyrimidine fraction
c
pmoCAB
Mean pyrimidines fraction
pmoCAB and of 1,000 random samples of n = 3
d
e
f
Percentage
T
T-mxa
T-xox
T-pmo
a
g
t
c
AT
GC
Purine
Pyrimidine
g
Mean nitrogen atoms per codon
pmoCAB and of 1,000 random samples of n = 3
Mean carbon atoms per codon
Carbon atoms per codon
Nitrogen atoms per codon
h
Mean oxygen atoms per codon
Hydrogen atoms per codon
Oxygen atoms per codon
i
MB_31_BR1
UCL = 0.06
r = 0.04
LCL = 0.01
UCL = 0.08
r = 0.05
LCL = 0.02
UCL = 0.05
r = 0.02
LCL = −0.01
UCL = 0.03
r = 0
LCL = −0.03
Nitrogen atoms per codon
Carbon atoms per codon
Oxygen atoms per codon
Hydrogen atoms per codon
mRNA abundance

**Figure S14.** Strategies for optimal transcription in *Methylobacter tundripaludum* 31/32. (a) Frequency of purine and pyrimidine content in all CDSs. (b) Normalized mRNA abundance and the pyrimidine content of the corresponding CDS of *M. tundripaludum* 31/32 growing under different conditions. (c) Mean number of pyrimidines per base pair of the *pmoCAB* CDSs and the mean number of pyrimidines per base pair of 1,000 random samples of three CDSs ($n = 3$). (d), (e), (f) Effects on the total ribonucleotide composition due to removing a set of transcribed CDSs from the transcriptome (T). (g), (h) Elemental composition of transcribed CDSs in *M. tundripaludum* 31/32. The main panel shows the number of element atoms per codon in each transcribed CDS with the dashed lines representing the mean of all transcribed CDSs in the genome. The top and right panels show the mean number of element atoms per codon of the *pmoCAB* transcribed CDSs and the mean number of element atoms per codon of 1,000 random samples of three transcribed CDSs. (i) Correlation between elemental composition and mRNA abundance. The Pearson correlation coefficient with 95% confidence levels is depicted in each panel.

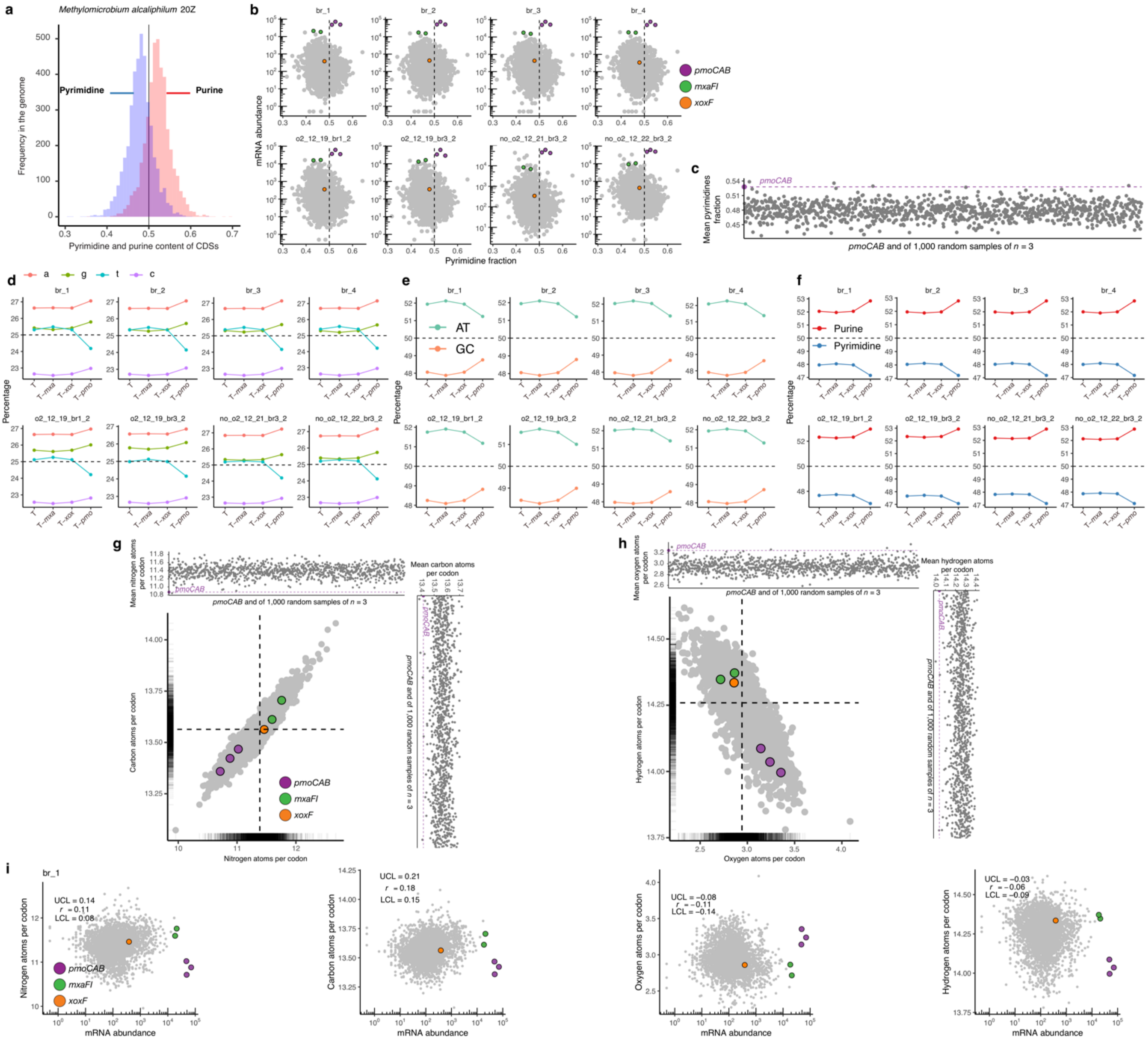

**Figure S15.** Strategies for optimal transcription in *Methylomicrobium alcaliphilum* 20Z. (a) Frequency of purine and pyrimidine content in all CDSs. (b) Normalized mRNA abundance and the pyrimidine content of the corresponding CDS of *M. alcaliphilum* 20Z growing under different conditions. (c) Mean number of pyrimidines per base pair of the *pmoCAB* CDSs and the mean number of pyrimidines per base pair of 1,000 random samples of three CDSs (*n* = 3). (d), (e)**,** (f) Effects on the total ribonucleotide composition due to removing a set of transcribed CDSs from the transcriptome (T). (g)**,** (h) Elemental composition of transcribed CDSs in *M. alcaliphilum* 20Z. The main panel shows the number of element atoms per codon in each transcribed CDS with the dashed lines representing the mean of all transcribed CDSs in the genome. The top and right panels show the mean number of element atoms per codon of the *pmoCAB* transcribed CDSs and the mean number of element atoms per codon of 1,000 random samples of three transcribed CDSs. (i) Correlation between elemental composition and mRNA abundance. The Pearson correlation coefficient with 95% confidence levels is depicted in each panel.

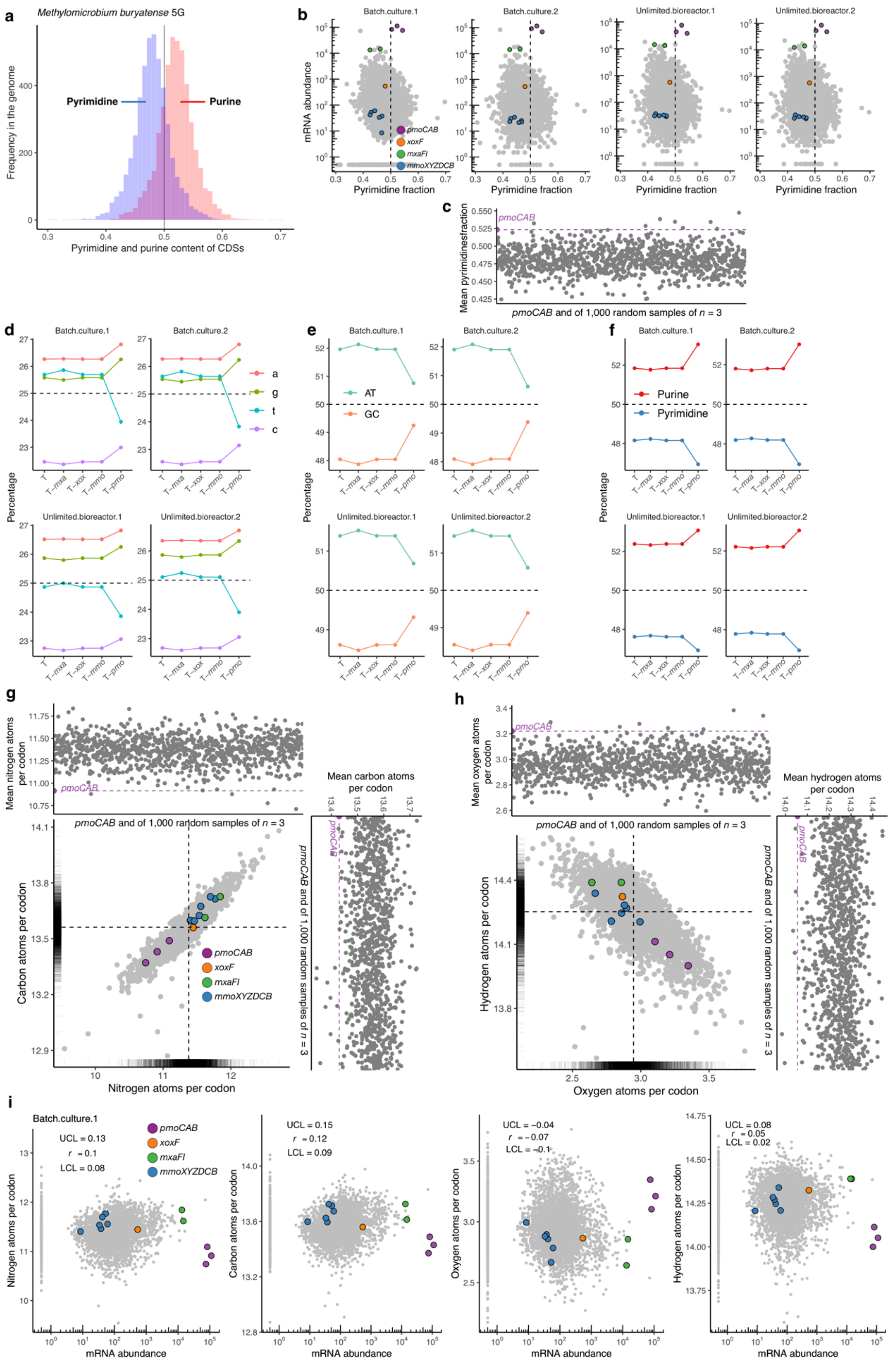

**Figure S16.** Strategies for optimal transcription in *Methylomicrobium buryatense* 5G. (a) Frequency of purine and pyrimidine content in all CDSs. (b) Normalized mRNA abundance and the pyrimidine content of the corresponding CDS of *M. buryatense*

5G growing under different conditions. (c) Mean number of pyrimidines per base pair of the *pmoCAB* CDSs and the mean number of pyrimidines per base pair of 1,000 random samples of three CDSs ($n$ = 3). (d), (e), (f) Effects on the total ribonucleotide composition due to removing a set of transcribed CDSs from the transcriptome (T). (g), (h) Elemental composition of transcribed CDSs in *M. buryatense* 5G. The main panel shows the number of element atoms per codon in each transcribed CDS with the dashed lines representing the mean of all transcribed CDSs in the genome. The top and right panels show the mean number of element atoms per codon of the *pmoCAB* transcribed CDSs and the mean number of element atoms per codon of 1,000 random samples of three transcribed CDSs. (i) Correlation between elemental composition and mRNA abundance. The Pearson correlation coefficient with 95% confidence levels is depicted in each panel.

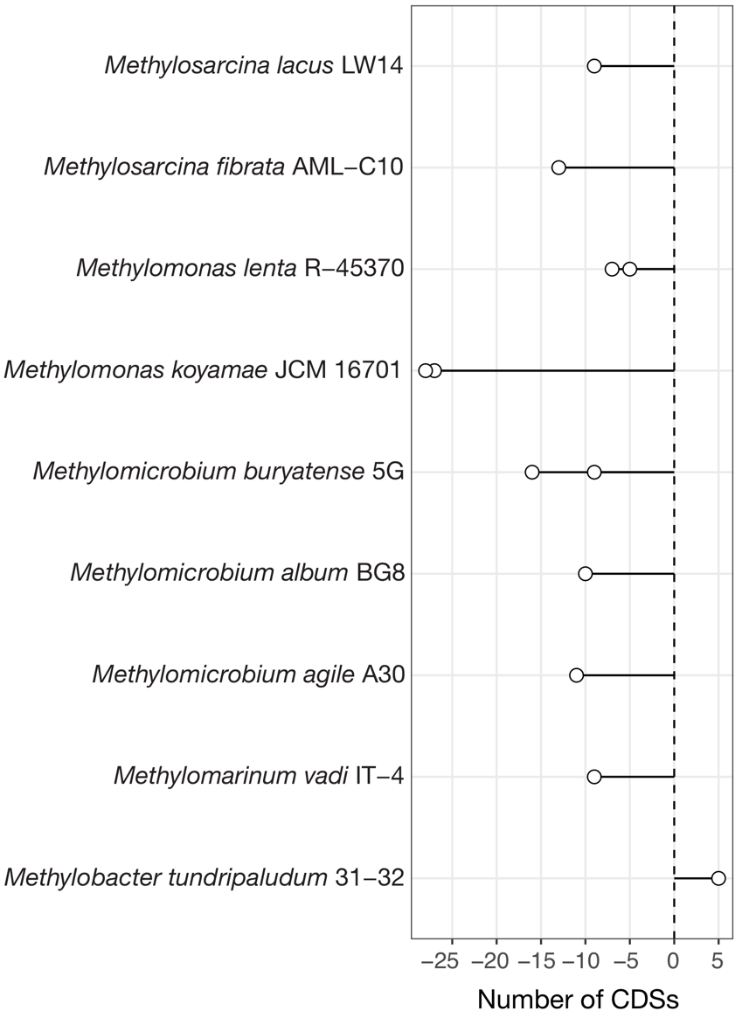

**Figure S17.** Location of the PAS domain proteins relative to the *pmoCAB* CDSs in different type Ia methanotrophs. Negative value in the *x*-axis indicates the number of CDSs upstream of the *pmoCAB* CDSs and positive value indicates downstream.

#### Additional file 2: Supplemental Table

**Table S1.** Detailed information and sources of the 67 genomes of methanotrophic bacteria including 49 isolate genomes and 18 MAGs used in this study.

#### References

1. R Core Team. R: A language and environment for statistical computing. R Foundation for Statistical Computing, Vienna, Austria. 2017.

2. RStudio Team. RStudio: Integrated development for R. RStudio, Inc., Boston, MA. 2015.

3. Wickham, H. ggplot2: elegant graphics for data analysis. Springer-Verlag New York. 2016.

12. Eren, M. A. et al. Anvi'o: an advanced analysis and visualization platform for 'omics data. PeerJ. 20153;e1319.

13. Wu, Y.-W., Simmons, B. A. & Singer, S. W. MaxBin 2.0: an automated binning algorithm to recover genomes from multiple metagenomic datasets. Bioinformatics. 2016;32:605–607.

14. Parks, D. H., Imelfort, M., Skennerton, C. T., Hugenholtz, P. & Tyson, G. W. CheckM: assessing the quality of microbial genomes recovered from isolates, single cells, and metagenomes. Genome Res. 2015;25:1043–1055.

15. Seemann, T. Prokka: rapid prokaryotic genome annotation. Bioinformatics. 2014;30:2068–2069.

16. Segata, N., Börnigen, D., Morgan, X. C. & Huttenhower, C. PhyloPhlAn is a new method for improved phylogenetic and taxonomic placement of microbes. Nat. Commun. 2013;4.

24. Becker, R. A., Wilks, A. R., Brownrigg, R., Minka, T. P. & Deckmyn, A. maps: draw geographical maps. R package version 3.3.0.2018.

25. Paradis, E., Claude, J. & Strimmer, K. APE: analyses of phylogenetics and evolution in R language. Bioinformatics 20, 289–290 (2004).

26. Hug, L. A. et al. A new view of the tree of life. Nat Microbiol. 2016;1:16048.

27. Yu, G. & Tsan-Yuk Lam, T. treeio: base classes and functions for phylogenetic tree input and output. R package version 1.2.2. 2018.

28. Yu, G., Smith, D. K., Zhu, H., Guan, Y. & Lam, T. ggtree: an R package for visualization and annotation of phylogenetic trees with their covariates and other associated data. Methods Ecol Evol. 2017;8:28–36.

29. Wilke, C. O. ggridges: ridgeline plots in ‘ggplot2’. R package version 0.5.0.2018.
